# Universal Cold RNA Phase Transitions

**DOI:** 10.1101/2024.03.22.586224

**Authors:** P. Rissone, A. Severino, I. Pastor, F. Ritort

## Abstract

RNA’s diversity of structures and functions impacts all life forms since *primordia*. We use calorimetric force spectroscopy to investigate RNA folding landscapes in previously unexplored low-temperature conditions. We find that Watson-Crick RNA hairpins, the most basic secondary structure elements, undergo a glass-like transition below T_G_ ∼ 20°C where the heat capacity abruptly changes and the RNA folds into a diversity of misfolded structures. We hypothesize that an altered RNA biochemistry, determined by sequence-independent ribose-water interactions, outweighs sequence-dependent base pairing. The ubiquitous ribose-water interactions lead to universal RNA phase transitions below T_G_, such as maximum stability at T_S_ ∼ 5°C where water density is maximum, and cold denaturation at T_C_ ∼ −50°C. RNA cold biochemistry may have a profound impact on RNA function and evolution.

---

Of similar chemical structure to DNA, the deoxyribose-ribose and thymine-to-uracil differences endow RNA with a rich phenomenology (1, 2, 3). RNA structures are stabilized by multiple interactions among nucleotides and water, often with the critical involvement of magnesium ions (4, 5, 6). Such interactions compete in RNA folding, producing a rugged folding energy landscape (FEL) with many local minima (7, 8). To be functional, RNAs fold into a native structure via intermediates and kinetic traps that slow down folding (9, 10). The roughness of RNA energy landscapes has been observed in ribozymes that exhibit conformational heterogeneity with functional interconverting structures (11, 12, 13) and misfolding (14, 15, 16, 17). Single-molecule methods have revealed a powerful approach to investigate these questions by monitoring the behavior of individual RNAs one at a time, using fluorescence probes (18, 19) and mechanical forces (20, 21). Previous studies have underlined the crucial role of RNA-water interactions at subzero temperatures in liquid environments (22, 23, 24) raising the question of the role of water in a cold RNA biochemistry. Here, we carry out RNA pulling experiments at low temperatures, showing that fully complementary RNA hairpins unexpectedly misfold below a characteristic glass-like transition temperature *T_G_* ∼ 20°C, adopting a diversity of compact folded structures. This phenomenon is observed in both monovalent and divalent salt conditions, indicating that magnesium-RNA binding is not essential for this to happen. Moreover, misfolding is not observed in DNA down to 5°C. These facts suggest that the folded RNA arrangements are stabilized by sequence-independent 2*^0^*-hydroxyl-water interactions that outweigh sequence-dependent base pairing. Cold RNA misfolding implies that the FEL is rugged with several minima that kinetically trap the RNA upon cooling, a characteristic feature of glassy matter (25). RNA folding in rugged energy landscapes is accompanied by a reduction of RNA’s configurational entropy. A quantitative descriptor of this reduction is the folding heat capacity change at constant pressure, Δ*C_p_*, directly related to the change in the number of degrees of freedom available to the RNA molecule. Despite its importance, Δ*C_p_* measurements in nucleic acids remain challenging (26, 27, 28). We carry out RNA pulling experiments at low temperatures and show that Δ*C_p_* abruptly changes at *T_G_* ∼ 20°C, a manifestation that the ubiquitous non-specific ribose-water interactions overtake the specific Watson-Crick base pairing at sufficiently low temperatures.

## RNA misfolds at low temperatures

We used a temperature-jump optical trap (Sec. 1, Methods) to unzip fully complementary Watson-Crick RNA hairpins featuring two 20bp stem sequences (H1 and H2) and loops of different sizes (*L* = 4, 8, 10, 12 nucleotides) and compositions (poly-A or poly-U) (Sec. 2, Methods). Pulling experiments were carried out in the temperature range 7 - 42°C at 4mM MgCl_2_ and 1M NaCl in a 100mM Tris-HCl buffer (pH 8.1). Figure 1A shows the temperature-dependence of the force-distance curves (FDCs) for the dodeca-A (12nt) loop hairpin sequence H1L12A at 4mM magnesium. At and above room temperature (*T* 2” 25°C), H1L12A unfolds at ∼ 20 - 25pN (blue force rips in dashed grey ellipse), and the rupture force distribution is unimodal (Fig. 1B, leftmost top panel at 25°C), indicating a single folded native state (N). At *T* ≤ 17°C, new unfolding events appear at higher forces (∼ 30 - 40pN, dashed black ellipse). The bimodal rupture force distribution (Fig. 1B, right top panels) shows the formation of an alternative misfolded structure (M) that remains kinetically stable over the experimental timescales. Below *T* = 10°C, the misfolded population shows *>* 50% occupancy. Analogous results were obtained with sodium ions (Fig. S2, Supp. Info). The formation of stable non-native structures for H1L12A is not predicted by algorithms such as Mfold (29), Vienna package (30), McGenus (31), pKiss (32), and Sfold (33). Furthermore, misfolding is not observed for the equivalent DNA hairpin sequence with deoxy-nucleotides (34). We refer to this phenomenon as cold RNA misfolding.

**Figure 1:**
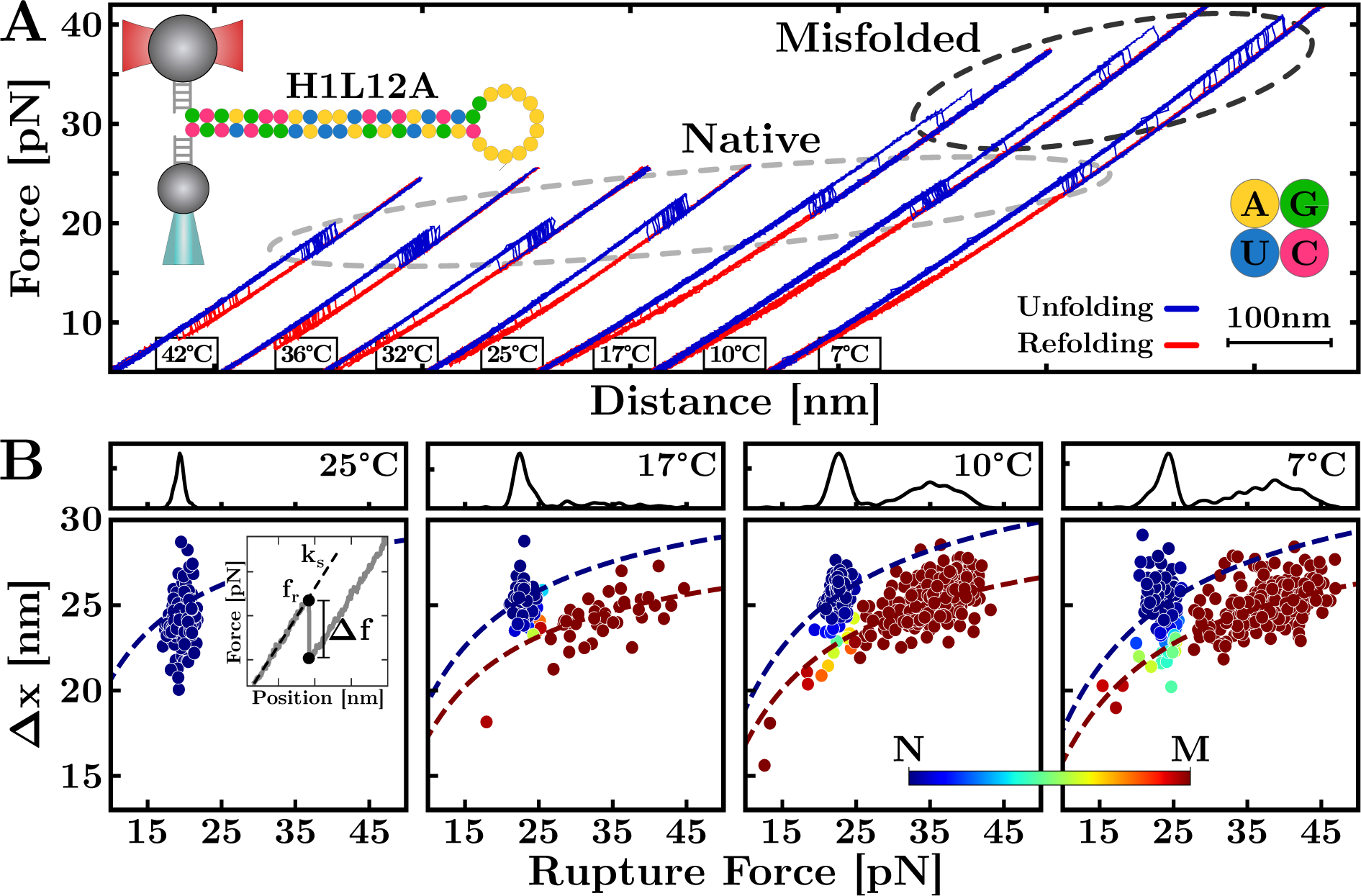
cold RNA misfolding. (**A**) Unfolding (blue) and refolding (red) FDCs from H1L12A unzipping experiments (top-left) at temperatures 7 - 42°C and 4mM MgCl_2_. The grey-dashed ellipse indicates native (N) unfolding events. Unexpected unfolding events from a misfolded (M) structure appear below 25°C (black-dashed ellipse) that become more frequent upon low-ering *T* from 17°C to 7°C. (**B**) Classification of N (blue dots) and M (red dots) rupture events at *T* ≤ 25°C and WLC fits for each state (dashed lines). The top panels show rupture force distributions at each *T*. The inset of the leftmost panel shows the parameters of rupture force events (see text).

Misfolding can be characterized by the size of the force rips at the unfolding events, which imply a change in the RNA molecular extension, Δ*x*. The value of Δ*x* is obtained as the ratio between the force drop Δ*f* and the slope *k*_s_ of the FDC measured at the rupture force *f_r_*, Δ*x* = Δ*f/k*_s_ (inset of left panel in Fig. 1B). Figure 1B shows Δ*x* versus *f_r_* for all rupture force events in H1L12A at four selected temperatures. Two clouds of points are visible below 25°C, evidencing two distinct folded states, the native (N, blue) and the misfolded (M, red). A Bayesian network model (Sec. 3, Methods) has been implemented to assign a probability of each data point belonging to N or M (color-graded bar in Fig. 1B). At a given force, the released number of nucleotides for N and M (*n_N_*, *n_M_*) is directly proportional to Δ*x* (Sec. S2, Supp. Info). To derive the values of *n_N_* and *n_M_*, a model of the elastic response of the single-stranded RNA (ssRNA) is required. We have fitted the datasets (Δ*x, f_r_*) for N and M to the worm-like chain (WLC) elastic model (Sec. 4, Methods) using the Bayesian network model, finding *n_N_*= 52(1) (blue dashed line) and *n_M_*= 46(1) (red dashed line) for the number of released nucleotides upon unfolding the N and M structures. Notice that *n_N_* matches the total number of nucleotides in H1L12A (40 in the stem plus 12 in the loop), while M features 6nt less than *n_N_*. These can be interpreted as remaining unpaired in M or that the 5*^0^* - 3*^0^* end-to-end distance in M has increased by ∼ 3nm, roughly corresponding to 6nt.

## RNA flexibility at low-***T*** promotes misfolding

To characterize the ssRNA elasticity, we show the force-extension curves versus the normalized ssRNA extension per base in Fig. 2A for H1L12A at different temperatures. Upon decreasing *T*, the range of forces and extensions becomes wider due to the higher unfolding and lower refolding forces. Moreover, a shoulder in the force-extension curve is visible below 32°C (see also Fig. S3, Supp. Info), indicating the formation of non-specific secondary structures. A similar phenomenon has been observed in ssDNA (35). The force-extension curves (triangles and circles in Fig. 2A) at each temperature were fitted to the WLC model, with persistence length *l_p_* and inter-phosphate distance *d_b_* as fitting parameters (Fig. 2B and Eq.(1) in Sec. 4, Methods). Only data above the shoulder has been used to fit the WLC (Sec. S1, Supp. Info). The values *l_p_* and *d_b_* show a linear *T*-dependence (red symbols in Fig. 2C) that has been used for a simultaneous fit of the ssRNA elasticity at all temperatures (blue lines in Fig. 2A). Over the studied temperature range, *l_p_* (Fig. 2C, left panel) increases with *T* by a factor of ∼ 2.5, whereas *d_b_* (Fig. 2C, right panel) decreases by only ∼ 20%. The increase of *l_p_* with *T* is an electrostatic effect (34) that facilitates the bending of ssRNA at the lowest temperatures, promoting base contacts and misfolding.

**Figure 2:**
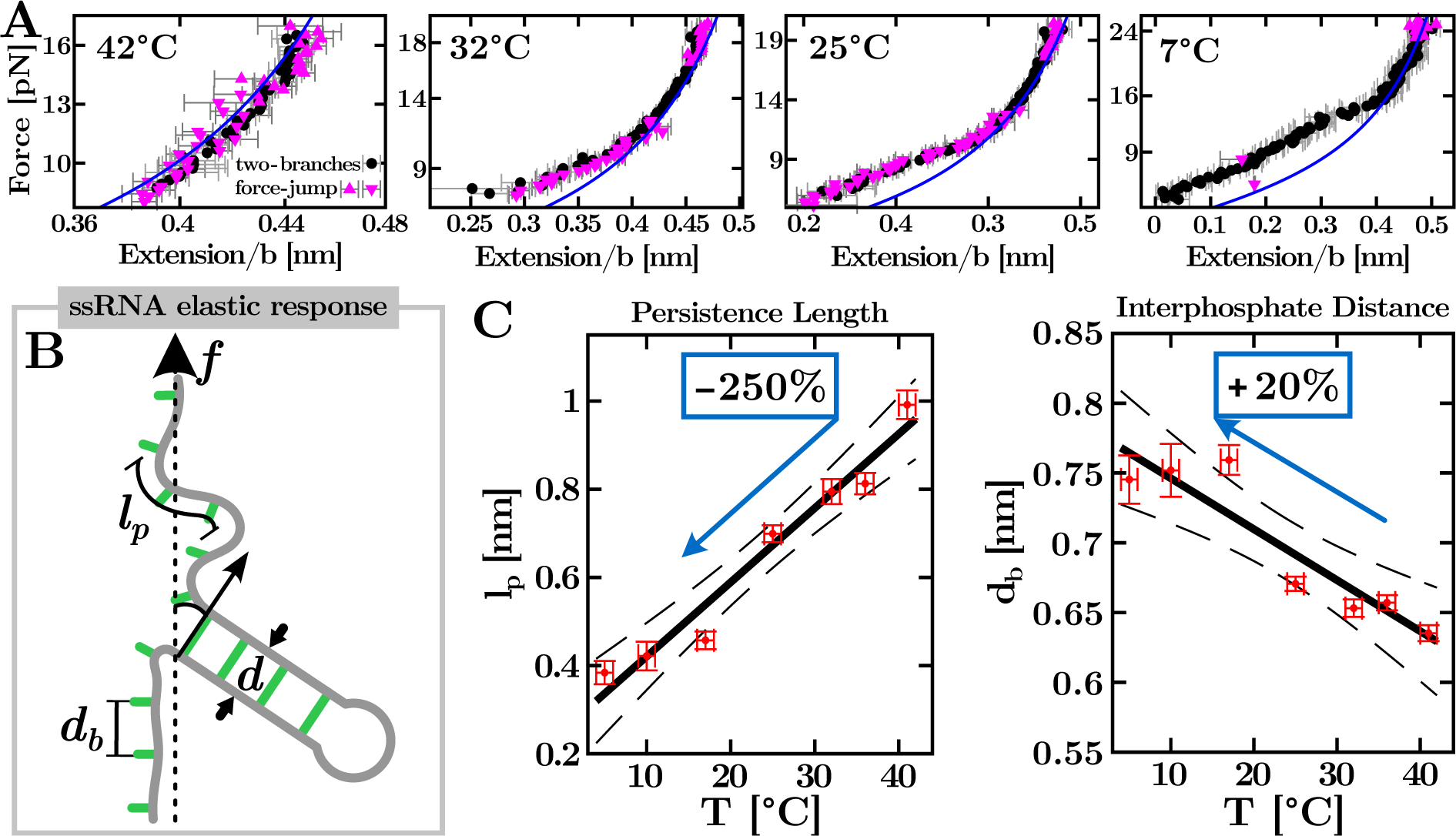
Temperature-dependent ssRNA elasticity. (**A**) Force versus the ssRNA extension per base at different temperatures. Two methods have been used to extract the ssRNA molecular extension: the force-jump (magenta triangles up – unfolding – and down – refolding –) and the two-branches method (black circles) (58, 59). Blue lines are the fits to the WLC in the high-force regime (see text). (**B**) Representation of the ssRNA elastic response according to the WLC model. The persistence length (*l_p_*) measures the polymer flexibility, and the interphosphate distance (*d_b_*) is the distance between contiguous bases. The computation of the total hairpin extension accounts for the contribution of the molecular diameter (*d*). (**C**) *T*-dependencies of *l_p_* (left) and *d_b_* (right). Linear fits (solid lines) with error limits (dashed lines) are also shown and give slopes equal to 0.17(2) Å/K for *l_p_* and −0.04(1) A Å/K for *d_b_*.

## Cold RNA misfolding is a universal sequence-independent phenomenon

The ubiquity of cold misfolding is due to the flexibility of the ssRNA rather than structural features such as stem sequence, loop size, and composition. To demonstrate this, we show results for another five hairpin sequences in Fig. 3A with different stem sequences and loop sizes. To assess the effect of loop size, three hairpins have the same stem as H1L12A but tetra-A, octa-A, and deca-A loops (H1L4A, H1L8A, H1L10A respectively). A fourth hairpin features a dodeca-U loop (H1L12U) to avoid base stacking in the dodeca-A loop of H1L12A. The fifth hairpin, H2L12A, has the same loop as H1L12A but features a different stem. Except for H1L4A, all hairpins misfold below *T* = 25°C, as shown by the emergence of unfolding events at forces above 30pN (blue rips in the black dashed ellipses in Fig. 3A) compared to the lower forces of the unfolding native events ∼ 20(grey dashed ellipses). Figure 3B shows the Bayesian-clustering classification of the different unfolding trajectories at 7°C and 25°C, in line with the results for H1L12A shown in Fig. 1B. The hairpin composition impacts misfolding; while H1L8A, H1L10A, and H1L12A show a single M at 7°C, H1L12U and H2L12A feature two distinct misfolded states at high (M_1_) and low (M_2_) forces (black dashed ellipses for H1L12U and H2L12A in Fig. 3A).

**Figure 3:**
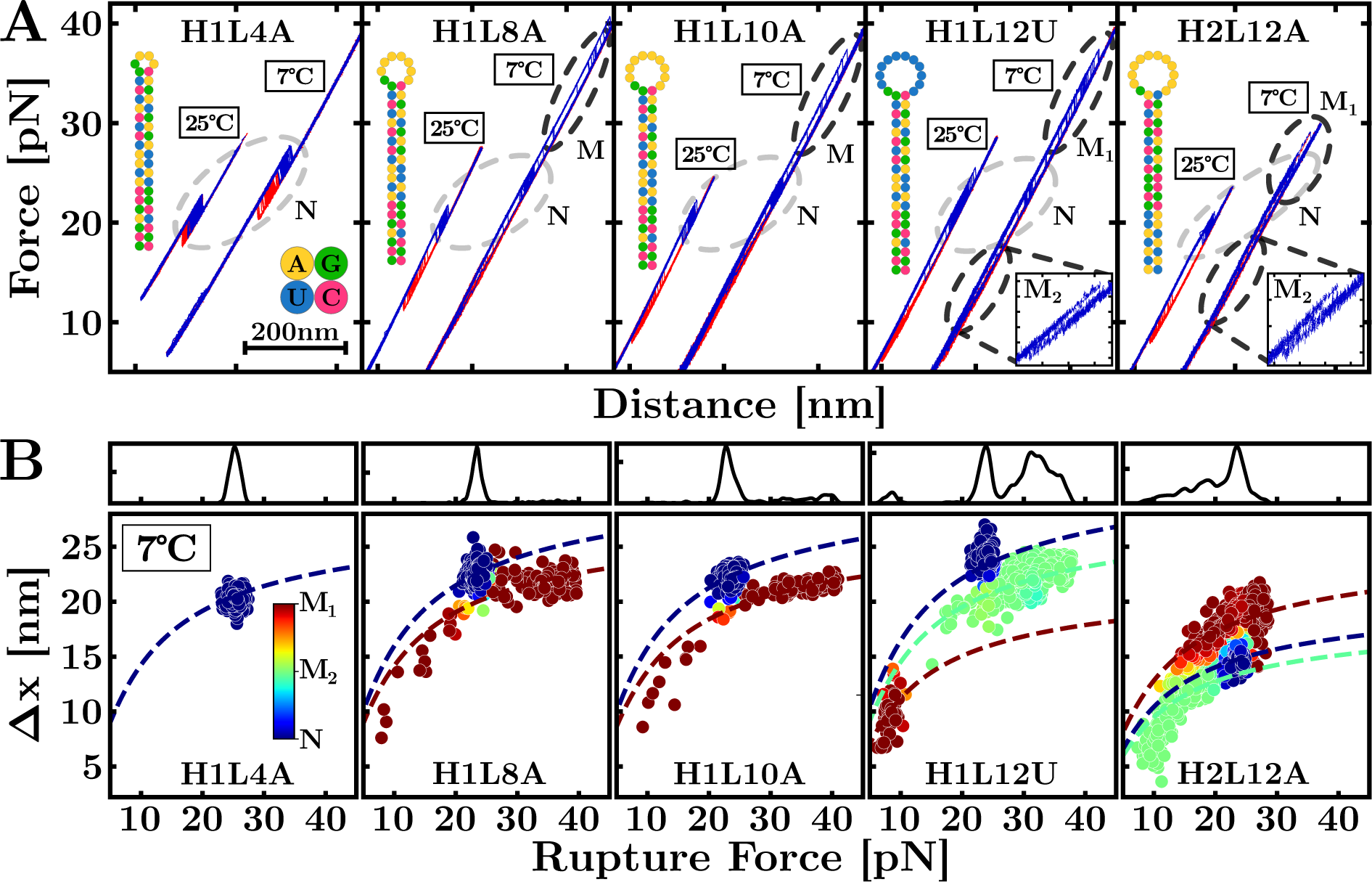
Universality of cold RNA misfolding. (**A**) Unfolding (blue) and refolding (red) FDCs of hairpins H1L4A, H1L8A, H1L10A, H1L12U, and H2L12A at 25°C and 7°C. Grey-dashed ellipses indicate native (N) unfolding events. Except for H1L4A, all RNAs show unfolding events from misfolded (M) structures at 7°C (black-dashed ellipses). Hairpins H1L12U and H2L12A (featuring a dodeca-U loop and a different stem sequence) show a second mis-folded structure at low forces (zoomed insets). Hairpin sequences are shown in each panel. (**B**) Bayesian classification of the unfolding events for the hairpins in panel (A) at *T* = 7°C. The dashed lines are the fits to the WLC for the different states. The top panels show the rupture force distributions.

The effect of the loop is to modulate the probability of formation of the native stem relative to other stable conformations. Indeed, H1L4A with a tetraloop has the largest stability among the studied RNAs (36), preventing misfolding down to 7°C (Fig. 3B). Misfolding prevalence increases with loop size due to the higher number of configurations and low entropic cost of bending the loop upon folding. The ssRNA elastic responses in H1L12A, H1L12U, and H2L12A show a systematic decrease of *l_p_* upon lowering *T* (Fig. S5, Supp. Info) and therefore an enhancement of misfolding due to the large flexibility of the ssRNA. Figure 4A shows the fraction of unfolding events at 7°C for all hairpin sequences for N (blue), M_1_ (red), and M_2_ (green). Starting from H1L4A, misfolding frequency increases with loop size, with the second misfolded state (M_2_) being observed for H1L12U and H2L12A within the limits of our analysis. Compared to the poly-A loop hairpins (Fig. S6, Supp. Info), the unstacked bases of the poly-U loop in H1L12U confer a larger *d_b_* and extension to the ssRNA (red dots in Fig. S5, Supp. Info). Elastic parameters for the family of dodecaloop hairpins are reported in Table S2, Supp. Info. The fact that hairpins containing poly-A and poly-U dodecaloops misfold at low temperatures demonstrates that stacking effects in the loop are nonessential to misfolding.

**Figure 4:**
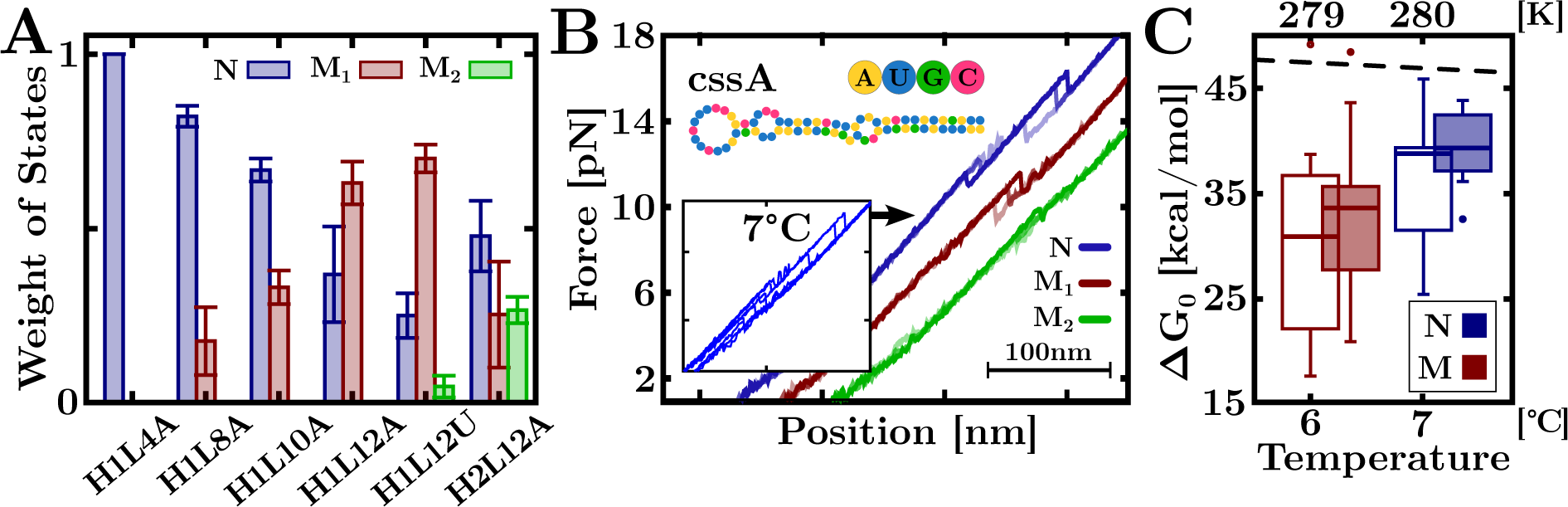
Features of cold RNA misfolding. (**A**) Frequency of N, M_1_, and M_2_ unfolding events for the different RNA hairpins at 7°C. (**B**) Unfolding FDCs of cssA RNA at 7°C and 4mM MgCl_2_ (inset) classified into native (N) and misfolded (M_1_, and M_2_) states. (**C**) Δ*G*_0_ values at 7°C in 4mM MgCl_2_ (solid boxes) and 400mM NaCl (empty boxes). Temperature axis in °C (bottom label) and K (top label). Box-and-whisker plots show the median (horizontal thick line), first and third quartiles (box), 10th and 90th percentiles (whiskers), and outliers (dots). The black dashed line is the Mfold prediction.

To further demonstrate the universality of cold RNA misfolding, we have pulled the mRNA of bacterial virulence protein CssA from *N. meningitidis*, an RNA thermometer that changes conformation above 37°C (37). Figure 4B shows several FDCs measured at 7°C and 4mM MgCl_2_ (inset), evidencing that the mRNA misfolds into two structures (M_1_, red; M_2_, green).

## RNA misfolds into stable and compact structures at low temperatures

The Bayesian analysis of the force rips has permitted us to classify the unfolding and refolding trajectories into two sets, *N* ⇌ *U* and *M* ⇌ *U* (Figs. 1B and 3B). We have applied the fluctuation theorem (38, 39) to each set of trajectories of H1L12A to determine the free energies of formation of N and M from the irreversible work (*W*) measurements at 7°C (Sec. 5, Methods and Sec. S4, Supp. Info). In Fig. 4C, we show Δ*G*_0_ estimates for N (blue) and M (red), finding Δ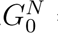 = 38(9) kcal/mol and Δ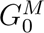 = 30(10) kcal/mol in 4mM MgCl_2_ (empty boxes). We have also measured Δ*G*_0_ at 1M NaCl and extrapolated it to 400mM NaCl, the equivalent concentration to 4mM MgCl_2_ according to the 100:1 salt rule (39). We obtain Δ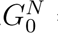 = 37(3) kcal/mol and Δ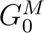 = 31(8) kcal/mol (filled boxes), in agreement with the magnesium data. Within the experimental uncertainties, Δ*G*_0_ for N is higher by ∼ 5 kcal/mol than for M, reflecting the higher stability of Watson-Crick base pairs in N. Notice that the Mfold prediction for N (Δ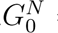 = 47 kcal/mol, black dashed line) overestimates Δ*G*_0_ by 10 kcal/mol.

We have also examined the distance between the folded and the transition state *x^‡^* in H1L12A to quantify the compactness of the folded structure. We have determined *x^‡^* from the rupture force variance *σ*^2^ using the Bell-Evans model, through the relation *σ*^2^ = 0.61(*k*_B_*T/x^‡^*)^2^ (Sec. S5, Supp. Info). We find that average rupture forces for N and M decrease linearly with *T*, whereas *σ*^2^ values are *T*-independent and considerably larger for M, *σ*^2^ ∼ 50*σ*^2^, giving 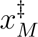= 0.7(4)nm and 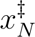 = 4.8(6)nm (Fig. S10, Supp. Info). Therefore, 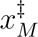 ⌧ 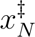 with M featuring a shorter *x^‡^* and a more compact structure than N.

## The RNA glassy transition

The ubiquity of the cold RNA misfolding phenomenon suggests that RNA experiences a glass transition below a characteristic temperature *T_G_* where the FEL develops multiple local minima. Figure 5A illustrates the effect of cooling on the FEL (40, 41). Above 25°C, the FEL has a unique minimum for the native structure N (red-colored landscape). The projection of the FEL along the molecular extension coordinate shows that N is separated from U by a transition state (TS) (top inset, red line). Upon cooling, the FEL becomes rougher with deeper valleys, promoting misfolding (green and blue colored landscapes). The distance from M to TS is shorter than from N to TS, reflecting that M is a compact structure (bottom inset, green and blue lines).

**Figure 5:**
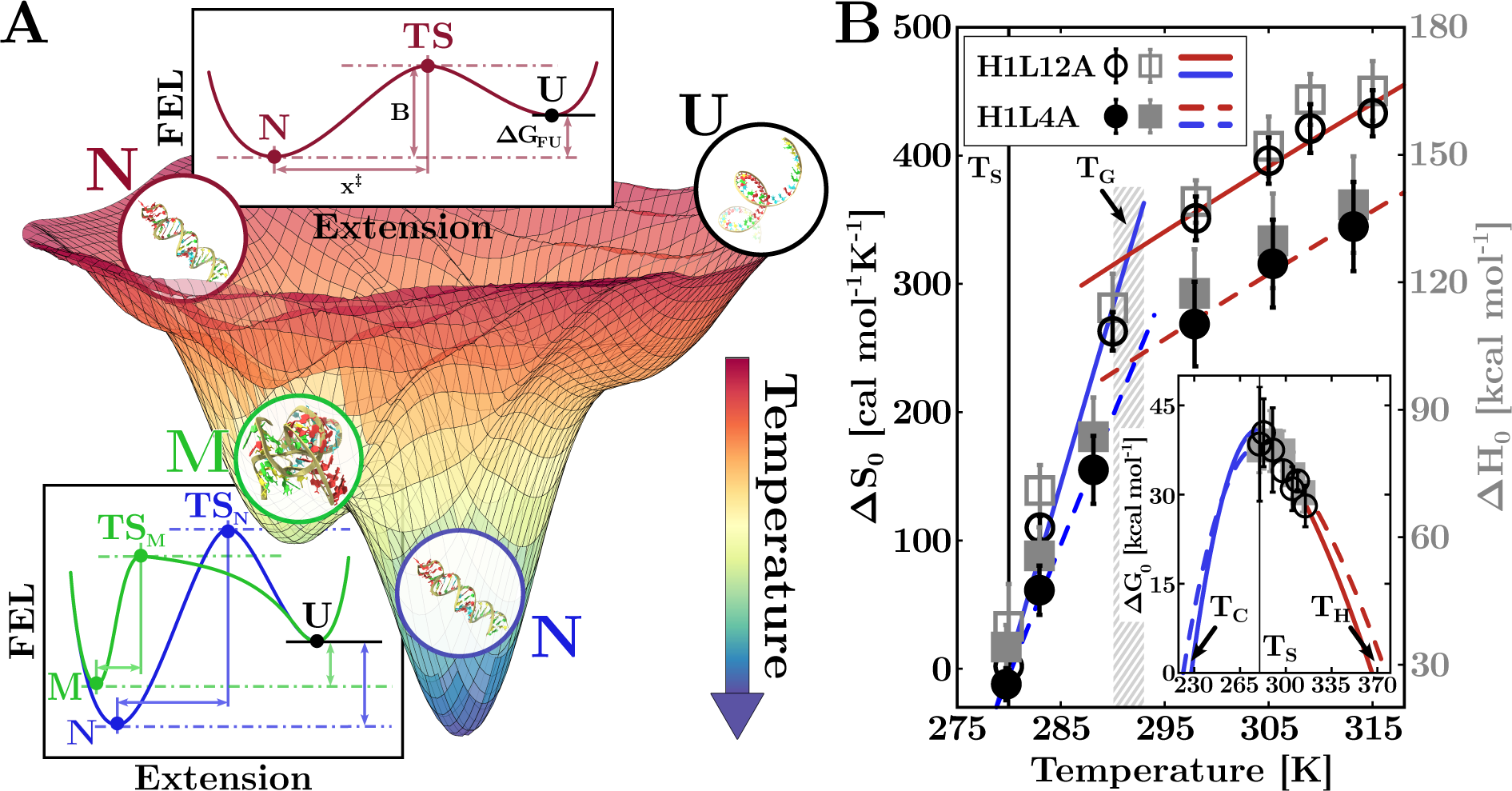
Cold RNA misfolding and phase transitions. (**A**) Illustration of a multi-colored free-energy landscape (FEL) at different temperatures. The temperature arrow indicates the tendency to explore low-lying energy states with the FEL becoming rougher upon cooling: from high (red) to intermediate (green) and low (blue) temperatures. Transition state (TS) distances are typically shorter for M than N, denoting disordered and compact misfolded structures. The encircled schematic folds are for illustration purposes. (**B**) Temperature-dependent entropy (black) and enthalpy (grey) of N for H1L12A (empty symbols) and H1L4A (full symbols). The results are reported in Table S5 and S6, Supp. Info. Fits to the entropy values in the hot (red) and cold (blue) regimes for H1L12A (solid lines) and H1L4A (dashed lines) are also shown. The transition between the two regimes occurs at *T_G_* ∼ 293K ∼ 20°C (dashed grey band) with a sudden change in Δ*C_p_*. **Inset.** Stability curves of H1L12A (empty black circles) and H1L4A (solid grey squares). Maximum stability is found at *T_S_* ∼ 278K ∼ 5°C (vertical black line) with melting temperatures at *T_H_* ∼ 370K ∼ 100°C (red lines). Extrapolations of Δ*G*_0_(*T*) in the cold regime predict cold denaturation at *T_C_* ∼ 220K ∼ −50°C for both hairpins (blue lines).

The glassy transition is accompanied by the sudden increase in the heat capacity change (Δ*C_p_*) between N and U below *T_G_* ∼ 20°C for H1L12A and H1L4A. Δ*C_p_* equals the temperature derivative of the folding enthalpy and entropy, Δ*C_p_* = *∂*Δ*H/∂T* = *T ∂*Δ*S/∂T* and can be determined from the slopes of Δ*H*(*T*) and Δ*S*(*T*) (Sec. 6, Methods and Sec. S8, Supp. Info). We observe two distinct regimes: above *T_G_* (hot, H) and below *T_G_* (cold, C). While 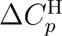 ∼ 1.5 · 10^3^ cal mol*^-^*^1^K*^-^*^1^ is similar for both H1L12A and H1L4A (parallel red lines in Fig. 5B), 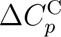 differs: 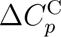 = 8(1)·10^3^ cal mol*^-^*^1^K*^-^*^1^ for H1L12A versus 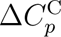 = 5.8(4)·10^3^ cal mol*^-^*^1^K*^-^*^1^ for H1L4A (unparallel blue lines in Fig. 5B) showing the dependence of 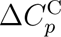 on loop size at low-*T*. Despite the different 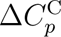 values, Δ*S*_0_ = 0 and stability (Δ*G*_0_) is maximum at *T_S_* = 5(2)°C (Fig. 5B, inset) for both H1L4A and H1L12A (vertical black lines in Fig. 5B, main and inset). Finally, the Δ*C*^C^ values predict cold denaturation at the same *T_C_* ∼ −50°C for both sequences. The agreement between the values of *T_G_*, *T_S_*, and *T_C_* suggests that cold RNA phase transitions are sequence-independent, occurring in narrow and well-defined temperature ranges for all RNAs.

## Discussion

Calorimetric force spectroscopy measurements on hairpin sequences of varying loop size, composition, and stem sequence show RNA misfolding at low-*T* in monovalent and divalent salt conditions. The phenomenon’s ubiquity leads us to hypothesize that non-specific ribose-water bridges overtake the preferential Watson-Crick base pairing of the native hairpin, forming compact structures at low temperatures. Cold misfolding is intrinsic to RNA, as it is not observed for the equivalent DNA hairpin sequences. In addition, magnesium ions are not crucial for it to happen, indicating the ancillary role of magnesium-mediated base-pairing interactions. Upon cooling, the diversity of RNA-water interactions promoted by the ribose increases the ruggedness of the folding energy landscape (FEL). Previous unzipping experiments of long (2kb) RNA hairpins at 25°C already identified stem-loops of ∼ 20 nucleotides as the *misfoldons* for RNA hybridization (39). The short RNA persistence length at low *T* (∼ 4Å ∼ at 10°C, Fig. 2C) facilitates non-native contacts between distant bases and the exploration of different configurations. Indeed, the higher flexibility of the U-loop in H1L12U enhances bending fluctuations and mis-folding compared to the stacked A-loop in H1L12A (Fig. 4A). Cold RNA misfolding has also been reported in NMR studies of the mRNA thermosensor that regulates the translation of the cold-shock protein CspA (42), aligning with the CssA results of Fig. 4B. Cold RNA misfolding should not be specific to force-pulling but also present in temperature-quenching experiments where the initial high-entropy random coil state further facilitates non-native contacts (43). We foresee that cold RNA misfolding might help to identify *misfoldon* motifs, contributing to developing rules for tertiary structure prediction (44, 45).

Most remarkable is the large 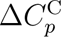 values for H1L12A and H1L4A below *T_G_* ∼ 20°C (293K), which are roughly 4-5 times the high-*T* value above *T_G_*, implying a large configurational entropy loss and a rougher FEL at low temperatures. The increase in Δ*C_p_* below *T_G_* (dashed grey band in Fig. 5B) is reminiscent of the glass transition predicted by statistical mod-els of RNA with quenched disorder (46, 47). As Δ*C_p_* = 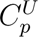 - 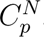, we attribute this change to the sudden reduction in 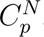 and the configurational entropy loss upon forming N (25). Both hairpins show maximum stability Δ*G*_0_ at *T_S_* ∼ 5°C (278K) where Δ*S*_0_ vanishes (Fig. 5B). The value of *T_S_* is close to the temperature where water density is maximum (4°C), with low-*T* extrapolations predicting cold denaturation at *T_C_* ∼ −50°C (220K) for both sequences. This result agrees with neutron scattering measurements of the temperature at which the RNA vibrational motion arrests, ∼ 220K (6, 24). We hypothesize that *T_S_* ∼ 5°C and *T_C_* ∼ −50°C mark the onset of universal phase transitions determined by the primary role of ribose-water interactions that are weakly modulated by RNA sequence, a result with implications for RNA condensates (48, 49) and RNA catalysis (50). The non-specificity of ribose-water interactions should lead to a much richer ensemble of RNA structures and conformational states and more error-prone RNA replication. Cold RNA could be relevant for extremophilic organisms, such as psychrophiles, which thrive in subzero temperatures (51). Finally, misfolding into compact and kinetically stable structures might help preserve RNAs in confined liquid environments such as porous rocks and interstitial brines in the permafrost of the arctic soil and celestial bodies (52, 53). This fact might have conferred an evolutionary advantage to RNA viruses for surviving during long periods (54) with implications on ecosystems due to the ongoing climate change (55). The ubiquitous sequence-independent ribose-water interactions at low temperatures frame a new paradigm for RNA self-assembly and catalysis in the cold. It is expected to impact RNA function profoundly, having potentially accelerated the evolution of a primordial RNA world (56, 57).

## Acknowledgments

We thank S. A. Woodson and C. Hyeon for a critical reading of the manuscript. **Funding**: P.R. was supported by the Angelo Della Riccia Foundation. I. P. and F.R. were supported by Spanish Research Council Grant PID2019-111148GB-I00 and the Institució Catalana de Recerca i Estudis Avançats (F. R., Academia Prize 2018). **Author contributions**: P.R., A.S., and I.P. carried out the experiments. P.R. and A.S. analyzed the data. I.P. and P.R. synthesized the molecules. I.P. and F.R. designed the research. P.R., A.S., and F.R. wrote the paper. **Competing interests**: The authors declare no competing financial interests. **Data and materials availability**: All data are available in the main text or the supplementary materials. This article has accompanying Supplementary Materials.

