## Supplementary Information for "Universal Cold RNA Phase Transitions"

### Supplementary Material for
#### Universal Cold RNA Phase Transitions

P. Rissone*, A. Severino*, I. Pastor, F. Ritort.
*Equally contributed authors.

**The PDF file includes:**

Materials and Methods
Supplementary Text
Supplementary Figs. S1 to S10
Supplementary Tables S1 to S6

**S2 Released nucleotides in unfolding events** **6**

**S3 Bayesian clustering** **7**

**S4 The H1L12A free-energies** **9**

**S5 Bell-Evans model** **10**

**S6 Continuous Effective Barrier Analysis** **11**

**S7 H1L12A thermodynamics for $\Delta C_p = 0$** **12**

**S8 H1L4A and H1L12A thermodynamics for $\Delta C_p \neq 0$** **13**

**Supplementary Figures** **14**

**Supplementary Tables** **24**

### Material and Methods

#### 1 Temperature-jump optical trap

We used a temperature-jump optical trap to perform unzipping experiments at different temperatures (60). Our setup adds to a MiniTweezers device (61) a heating laser of wavelength $\lambda = 1435$nm to change the temperature inside the microfluidics chamber. The latter is designed to damp convection effects caused by the laser non-uniform temperature, which may produce a hydrodynamics flow between medium regions (water) at different $T$. The heating laser allows for increasing the temperature by discrete amounts of $\Delta T \sim +2.5$°C up to a maximum of $\sim +30$°C with respect to the environment temperature, $T_0$. Operating the instrument in an icebox cooled down at a constant $T_0 \sim 5$°C, and outside the box at ambient temperature (25°C), we carried out experiments in the $T$ range $[7, 42]$°C.

In a pulling experiment, the molecule is tethered between two polystyrene beads through specific interactions with the molecular ends (62). One end is labeled with a digoxigenin (DIG) tail and binds with an anti-DIG coated bead (AD) of radius $3\mu$m. The other end is labeled with biotin (BIO) and binds with a streptavidin-coated bead (SA) of radius $2\mu$m. The SA bead is immobilized by air suction at the tip of a glass micropipette, while the AD bead is optically trapped. The unfolding process is carried out by moving the optical trap between two fixed positions: the molecule starts in the folded state, and the trap-pipette distance ($\lambda$) is increased until the hairpin switches to the unfolded conformation. Then, the refolding protocol starts, and $\lambda$ is decreased until the molecule switches back to the folded state.

The unzipping experiments were performed at two different salt conditions: 4mM $MgCl_2$ (divalent salt) and 1M NaCl (monovalent salt). Both buffers have been prepared by adding the salt (divalent or monovalent) to a background of 100mM Tris-HCl (pH $8.1$) and $0.01\%$ $NaN_3$. The NaCl buffer also contains 1mM EDTA. The pulling protocols have been carried out at a constant pulling speed, $v = 100$nm/s. We sampled 5-6 different molecules for each hairpin and at each temperature, collecting at least $\sim 200$ unfolding-folding trajectories per molecule.

#### 2 RNA synthesis

We synthesized six different RNA molecules made of a 20bp fully complementary Watson-Crick stem, ending with loops of different lengths ($L = 4, 8, 10, 12$ nucleotides) and compositions (poly-A or poly-U). The hairpins are flanked by long hybrid DNA/RNA handles ($\sim 500$bp). Further details about the sequences are given Fig. S1 and Table S1, Supp. Info.

The RNA hairpins have been synthesized using the steps in Ref.(63). First, the DNA template (Merck, Township, NJ, USA) of the RNA is inserted into plasmid pBR322 (New England Biolabs, NEB, Ipswich, MA, USA) between the HindIII and EcoRI restriction sites and cloned into the *E. coli* ultra-competent cells XL10-GOLD (Quickchange II XL site-directed mutagenesis kit). Second, the DNA template is amplified by PCR (KOD polymerase, Merck) using T7 promoters. The RNA is obtained by *in-vitro* RNA transcription (T7 megascript, Merck) of the DNA containing the RNA sequence flanked by an extra 527 and 599 bases at the 3′-end and 5′-end, respectively, for the hybrid DNA-RNA handles. Finally, labeled biotin (5′-end) and digoxigenin (3′-end) DNA handles, complementary to the RNA handles, are hybridized to get the final construct.

#### 3 Bayesian clustering

We use a mixture hierarchical Bayesian model (probabilistic graph network) to classify unfolding events as either emanating from a native or a misfolded initial folded state. The model is a soft classifier, giving each trace a probability (score) to belong to a given state. The model is described in Sec. S3, Supp. Info.

#### 4 ssRNA elastic model

The ssRNA elastic response has been modeled according to the worm-like chain (WLC), which reads

$$f(x) = \frac{k_{\mathrm{B}}T}{4l_p}\left[\left(1-\frac{x}{nd_b}\right)^{-2} - 1 + 4\frac{x}{nd_b}\right], \tag{1}$$

where $l_p$ is the persistence length, $d_b$ is the interphosphate distance and $n$ is the number of bases of the ssRNA. More details on the WLC model and the fitting method used to derive its parameters can be found in Sec. S1, Supp. Info.

#### 5 Free energy determination

Given a molecular state, $\Delta G_0(N)$ is the hybridization free energy of the $N$ base pairs of the folded structure when no external force is applied ($f = 0$). $\Delta G_0(N)$ is obtained from the free energy difference, $\Delta G_\lambda$, between a minimum ($\lambda_{\mathrm{min}}$) and a maximum ($\lambda_{\mathrm{max}}$) optical-trap positions where the molecule is folded and unfolded, respectively. Thus, one can write

$$\Delta G(\lambda) = \Delta G_0(N) + \Delta G_{\mathrm{el}}(\lambda), \tag{2}$$

where $\Delta G_{\mathrm{el}}(\lambda)$ is the elastic energy upon stretching the ssRNA between $\lambda_{\mathrm{min}}$ and $\lambda_{\mathrm{max}}$. The latter term can be computed by integrating the WLC (Eq.(1)). As unzipping experiments are performed by controlling the optical-trap position (not the force), this requires inverting Eq.(1) (Sec. S1, Supp. Info.).

We used the fluctuation theorem (64) (FT) to extract $\Delta G(\lambda)$ from irreversible work ($W$) measurements. This is computed by integrating the FDC between $\lambda_{\mathrm{min}}$ and $\lambda_{\mathrm{max}}$, $W = \int_{\lambda_{\mathrm{min}}}^{\lambda_{\mathrm{max}}} f d\lambda$ (inset in Fig. S8). Given the the forward ($P_{\mathrm{F}}(W)$) and reverse ($P_{\mathrm{R}}(W)$) work distributions, the FT reads

$$\frac{P_{\mathrm{F}}(W)}{P_{\mathrm{R}}(-W)} = \exp\left(\frac{W-\Delta G(\lambda)}{k_{\mathrm{B}}T}\right), \tag{3}$$

where the minus sign of $P_{\rm R}(-W)$ is due to the fact that $W < 0$ in the reverse process. When the work distributions cross, i.e. $P_{\rm F}(W) = P_{\rm R}(-W)$, Eq.(3) gives $W = \Delta G(\lambda)$. Let us notice that the FT can only be applied to obtain free-energy differences between states sampled under equilibrium conditions. However, pulling experiments at low $T$ are carried out under partial equilibrium conditions, with misfolding being a kinetic state. It is possible to extend the FT to our case by adding to $\Delta G(\lambda)$ from Eq.(3) the correction term $k_{\rm B}\log\left(\phi^i_{\rm F}/\phi^i_{\rm R}\right)$, where $\phi_{\rm U(R)}$ is the fraction of forward (reverse) trajectories of state $i = {\rm N}, {\rm M}$ (38). Given $\Delta G(\lambda)$, the free energy at zero force, $\Delta G_0(N)$, is computed from Eq.(2) by subtracting the energy contributions of stretching the ssRNA, the hybrid DNA/RNA handles, and the bead in the trap. The first two terms are obtained by integrating the WLC in Eq.(1)), while the latter is modeled as a Hookean spring of energy $\Delta G_b(x) = 1/2k_{\rm b}x^2$, where $k_{\rm b}$ is the stiffness of the optical trap.

### 6 Derivation of the heat capacity change

To derive $\Delta C_p$, we have measured the enthalpy $\Delta H_0$ and entropy $\Delta S_0$ of N at different $T$'s for H1L12A and H1L4A. $\Delta S_0$ is obtained from the extended form of the Clausius-Clapeyron equation in a force (61), while $\Delta H_0 = \Delta G_0 + T\Delta S_0$. Both $\Delta H_0$ and $\Delta S_0$ are temperature dependent, with a finite $\Delta C_p$ (Sec. S8, Supp. Info). This has been obtained by fitting the $T$-dependent entropies to the thermodynamic relation $\Delta S_0(T) = \Delta S_m + \Delta C_p \log\left(T/T_m\right)$, where $T_m$ is the reference temperature and $\Delta S_m$ is the entropy at $T = T_m$.

### Supplementary Text

### S1 Worm-like chain model

#### 1 Explicit inversion

We describe the ssRNA elastic response according to the worm-like chain (WLC) model (65, 66), which reads

$$f_{\mathrm{WLC}}(x) = \frac{k_{\mathrm{B}}T}{4l_p}\left[\left(1-\frac{x}{nd_b}\right)^{-2} - 1 + 4\frac{x}{nd_b}\right], \tag{S1}$$

where $l_p$, $d_b$, and $n$ are the persistence length, interphosphate distance, and number of monomers in the ssRNA, respectively. Note that the WLC model expresses the force as a function of the extension and can be inverted (67) to obtain the extension per monomer $z \equiv x/nd_b$ as a function of the force:

$$z(f) = f^{-1}_{\mathrm{WLC}}(f)\,. \tag{S2}$$

The inverted form of the WLC, $z(f)$, speeds up data analysis and is implemented in the JAGS library for the Bayesian classification (Sec. S3, Methods).

#### 2 Multi-$T$ elastic response fit

We fit the temperature dependence of the elastic parameters ($l_p$, $d_b$) to the WLC model, assuming they are linearly dependent on temperature. The assumption is supported by available experimental evidence (Fig. 2C, main text). Therefore, we use the fitting expressions:

$$l_p = l_p(T) = a_1 T + b_1 \tag{S3a}$$
$$d_b = d_b(T) = a_2 T + b_2\,. \tag{S3b}$$

The result of this multi-$T$ fitting procedure is shown in Fig. S4, Supp. Info. Notice that to avoid the secondary structure plateau (which cannot be described by the WLC model), only data points in the high force range of the elastic response above the shoulder in the data have been used.

### S2 Released nucleotides in unfolding events

In an unfolding event, the extension of the ssRNA released in the transition from the folded to the unfolded state, $x_r(f)$, can be obtained from the experimental FDCs through the relation

$$x_r(f^U) = \frac{\Delta f}{k^F_{\mathrm{eff}}} + x_d(f^U), \tag{S4}$$

where $\Delta f = f^F - f^U$ is the force difference upon unzipping between the force in the folded branch $F$ ($f^F$) and in the unfolded branch $U$ ($f^U$), $k_{\text{eff}}^F$ is the effective stiffness in the folded branch, i.e. the slope of the FDC before unfolding, and $x_d$ is the diameter of the folded structure projected along the pulling axis. We used the value of $x_r$ determined from Eq.(S4) to asses whether the unfolding events experimentally observed originate from the native state (hairpin) or misfolded state. Given the number of nucleotides in the folded structure, $n$, the following relation holds:

$$x_r(f^U) = n \cdot d_b \cdot f_{\text{WLC}}^{-1}(f^U)\,. \tag{S5}$$

Therefore, different states characterized by different $n$ give different ($f^U$, $x_r(f^U)$) distributions, as shown in Fig. 1B and 3B, of the main text.

The advantage of Eq.(S5) is two-fold. First, by assuming that the WLC parameters $l_p$, $d_b$ (see Sec. S1) are known, the equation can be applied to infer the number of monomers $n$ in the folded structure, as is done in the Bayesian hierarchical model presented in Sec. S3. By determining $n$, we can also distinguish whether the RNA has folded into the native state or a misfolded state. Second, assuming $n$ to be known, the equation can be used to determine the value of the WLC parameters $l_p$, $d_p$ with a least squares fitting method. For H1L12A, the native state has $n = 52$, permitting us to derive the ssRNA elastic parameters.

### S3 Bayesian clustering

To model the unzipping experiments of RNA at low temperatures, we used a Bayesian network approach (mixture hierarchical Bayesian model). This has two advantages. First, using latent state variables in the model gives posterior distributions for the state of each data point, allowing a probabilistic soft clustering of each unfolding trace, i.e. the probability of the RNA being misfolded or native is assigned to each point. Second, using appropriate likelihood functions in the model gives a range of useful physical parameters, such as the mode and scale parameters of the rupture force distribution of each state. These parameters are related to the force average and variance. The latter gives us estimates of the distance to the transition state, $x^{\ddagger}$, (see Sec. S5), and the weight of each state, native and misfolded, in the total population.

We recall that Bayesian network models posit that the prior distributions of the parameters to be estimated are known. Similarly, the likelihood function to observe each data point *given* these prior parameters is known. The estimation of the model parameters is then obtained by computing the posterior distribution of the model, given by the Bayes theorem:

$$\mathit{Posterior} \propto \mathit{Likelihood} \times \mathit{Prior}\,, \tag{S6}$$

which is often done in practice with Monte Carlo methods.

In RNA unzipping experiments, the model data points are the pairs ($f$, $x_r$) that characterize the rupture force and released extension of each unfolding event in the forward unzipping process. The model core idea is that the extension ($x_r$) released in an unfolding event depends both on the initial folded state of the molecule (through the number of released monomers $n$, see Sec. S2) and the rupture force, $f$, since force distributions are state-dependent, Sec. S5. In practice, we use Eq.(S5) assuming that it is valid down to the presence of experimental noise, characterized as the difference between the r.h.s and l.h.s of Eq.(S5) and which we posit to be Laplace distributed around $0$ with precision $t$ (we comment further on this distribution choice at the end of the section). Therefore, for each data point ($f_i$, $x_{r,i}$), we have:

$$x_{r,i} - n_{z_i} \cdot d_B \cdot f_{\mathrm{WLC}}^{-1}(f_i) \sim \mathrm{Laplace}(0, t)\,, \tag{S7}$$

where the dependence on the number of monomers released in an unfolding event, $n$, is introduced through the use of the so-called latent variable $z_i$, which captures the initial state of a trajectory for each data point $i = 1, .., N$. Here, we use the shorthand notation $z_i = 1$ for the native state and $z_i = 2$ for the misfolded state.

The second core idea of the Bayesian classification consists of explicitly modeling the state dependency of the rupture force distribution. We posit that the parameters underpinning the rupture force distribution depend on the latent variable $z_i$. More specifically, we assume that rupture forces are Gompertz distributed with mode $M$ and scale $1/s$, and we have therefore set $M \equiv M_{z_i}$ and $s \equiv s_{z_i}$ with different values for the native/misfolded states. The overall likelihood of observing an experimental point $(f_i, x_{r,i})$ is obtained by putting all these elements together:

$$\mathit{Likelihood} = p\left(\overline{f_{\mathrm{WLC}}^{-1}}(f_i, n_{z_i}) - x_{r,i} |\, 0, t\right) \times p\left(f_i |\, M_{z_i}, s_{z_i}\right) \times p\left(z_i |\, \vec{w}\right)\,. \tag{S8}$$

The first term on the r.h.s is based on Eq.(S7) as described above, with the shortened notation $\overline{f_{\mathrm{WLC}}^{-1}}(f_i, n_{z_i}) \equiv n_{z_i} \cdot d_B \cdot f_{\mathrm{WLC}}^{-1}(f_i)$. The second term is given by the Gompertz likelihood mentioned above. The third term, $p\left(z_i|\vec{w}\right)$ represents the likelihood of the latent variable $z_i$ given a weight vector $\vec{w} = (w_1, w_2)$ whose components give the average occupancy of each state. We use for $z_i$ the standard conjugate pair of a Categorical distribution for the likelihood $p$ combined with a Dirichlet prior for $\vec{w}$.

The formal specification of the model can then be finally completed by defining appropriate priors for the parameters we want to infer, namely $n_1$, $n_2$, $M_1$, $M_2$, $s_1$, $s_2$, $t$, and $\vec{w}$. As already mentioned, we use for $\vec{w}$ a Dirichlet prior and parameterize both $t$ and $s_1$,$s_2$ with gamma priors. Finally, we take normal priors for $n_1$,$n_2$, and Laplace priors for $M_1$,$M_2$. The overall prior is then given by

$$\begin{aligned}\mathit{Prior} =& p(n_{z_i} |\, \mu_{z_i}, \nu_{z_i}) \times p\left(M_{z_i} |\, \tilde{\mu}_{z_i}, \tilde{\tau}_{z_i}\right) \times p\left(s_{z_i} |\, \tilde{\phi}_{z_i}, \tilde{\omega}_{z_i}\right) \times \\ & \times p(\vec{w} |\, \vec{\alpha}) \times p(t |\, \phi, \omega)\,,\end{aligned} \tag{S9}$$

where the model hyper-parameters are made explicit with $z_i = 1, 2$. Hyper-parameters are given by the Greek variables $\mu_1$, $\mu_2$, $\nu_1$, $\nu_2$, $\tilde{\mu}_1$, $\tilde{\mu}_2$, $\tilde{\tau}_1$, $\tilde{\tau}_2$, $\tilde{\phi}_1$, $\tilde{\phi}_2$, $\tilde{\omega}_1$, $\tilde{\omega}_2$, $\alpha$, $\phi$ and $\omega$. We emphasize that while different valid choices of priors could be made, all the priors chosen here were purposefully parameterized to be very flat in order to minimally constrain the posterior space.

Given the likelihood function and our choice of priors, we use Bayes theorem to compute the posterior distribution of the parameters we want to infer:

$$p(\{z_i\}_{i=1}^{N}; n_{1,2}; w_{1,2}; M_{1,2}; s_{1,2}; \sigma) \propto \mathit{Likelihood} \times \mathit{Prior}\,, \tag{S10}$$

where we defined for convenience $\sigma = 1/t$, the inverse of the precision $t$. The model with all its priors, likelihood, and variables is schematically summarized in Fig. S7, Supp. Info.

We used the R library RJAGS (68) to set up the Bayesian network. Posterior distributions were obtained by running at least three Monte Carlo Markov Chains (MCMC) using the RJAGS library, with a burn-in of 1000 iterations, followed by 5000 iterations. We ran the usual convergence and diagnostics test for MCMCs (Gelmann, chain intercorrelation coefficient) and visually inspected the MCMC noise term to confirm that our simulations converged. We always took the median of the posterior distribution of interest for point estimates (e.g., $n_1$, $n_2$). We give some additional details on other important aspects of the fitting procedure:

- The rupture forces are modeled as Gompertz-distributed. This is usually a good approximation in practice and even true in the BE model, Sec. S5. Each rupture force distribution (misfolded/native) is then parametrized by a different mode $M_{z_i}$ and scale parameter $1/s_{z_i}$. Note, however, that JAGS/RJAGS does not offer a Gompertz likelihood function by default. Therefore, we need to input the likelihood manually, using Eq.(S15) and the zero trick.

- When designing the model, we initially modeled the noise term in the l.h.s of Eq.(S7) with a more standard Gaussian likelihood. However, we quickly realized that some experimental points could feature large deviations between $x_i$ and $\overline{f_{\text{WLC}}^{-1}}(f_i, n_{z_i})$, deviations which skew/bias the model when assuming normality and lead to overall poor convergence performance of the Monte Carlo Markov Chain (MCMC) simulation. Hence, we choose to use a more robust Laplace likelihood, which is more accommodating when a few large outliers are present. This considerably improved the model's stability. Moreover, the Deviance Information Criterion (DIC) score of the model with Laplace likelihood was lower than with a Gaussian model, giving further confidence in this choice.

### S4 The H1L12A free-energies

Mechanical work measurements were extracted from unzipping data as described in Sec. 5, Methods. The inset of Fig. S8A illustrates the work measured between two fixed positions (vertical lines) for the unfolding (red) and refolding (blue) FDCs in a given $N \rightleftharpoons U$ cycle. Let $P_{\rightarrow}(W), P_{\leftarrow}(-W)$ and $\Delta G$ denote the work distributions and free energy difference between N or M and U. In Fig. S8A we show $P_{\rightarrow}(W)$ and $P_{\leftarrow}(-W)$ for H1L12A above room temperature, where only N is observed. For the work, $W$, we have subtracted the energy contributions of stretching the ssRNA, the hybrid DNA/RNA handles, and the bead in the trap (Sec. 5, Methods). The free energy at zero force, $\Delta G_0$, has been obtained by applying statistical approaches such as the Bennett acceptance ratio (BAR) method (39). Additionally, we have also determined $\Delta G_0$ using a diffusive kinetics model for the unfolding reaction, the so-called continuous effective barrier analysis (CEBA) (69) (see Sec. S6). In Fig. S9 (right panel), we show $\Delta G_0$ values obtained with BAR (blue circles) and CEBA (red circles) above 25°C (Table S3) finding compatible results. The value of $\Delta G_0$ agrees with the Mfold prediction (29) at 37°C (black triangles). However, a large discrepancy is observed for the enthalpy and entropy values if we assume $\Delta C_p = 0$, suggesting a non-zero $\Delta C_p$ (Sec. S7 and main text).

Using the BAR method, we have also determined $\Delta G_0$ for N and M at 7°C. In Fig. S8B, we show work distributions for N (top) and M (bottom) along with $\Delta G_0$ estimates (grey bands), finding $\Delta G_0^N = 68(16)\, k_\mathrm{B}T$ (38(9) kcal/mol) and $\Delta G_0^M = 54(18)\, k_\mathrm{B}T$ (30(10) kcal/mol) in 4mM $MgCl_2$. We have also measured $\Delta G_0^M$ at 1M NaCl and extrapolated it to 400mM NaCl, the equivalent concentration to 4mM $MgCl_2$ according to the 100:1 salt rule (39). We obtain $\Delta G_0^N = 37(3)$ kcal/mol and $\Delta G_0^M = 31(8)$ kcal/mol in 400mM NaCl in agreement with the magnesium data. Figure Fig. S9 (left panel) shows $\Delta G_0$ for 4mM $MgCl_2$ (filled boxes) and 400mM NaCl (empty boxes). These values agree with a linear extrapolation from high temperatures to 7°C (blue and red lines, right panel). In contrast, the Mfold prediction ($\Delta G_0^N =$ 47 kcal/mol, black dashed line) overestimates $\Delta G_0$ by 10 kcal/mol.

### S5 Bell-Evans model

According to the BE model (70, 71), the unfolding and folding kinetic rates between the folded ($F$) state and the unfolded ($U$) state, can be written as

$$k_{F\to U}(f) = k_0 \exp\left(-\frac{B_0 - f x^\ddagger}{k_\mathrm{B}T}\right) \tag{S11a}$$

$$k_{U\to F}(f) = k_0 \exp\left(-\frac{B_0 - \Delta G_{FU} + f(x_U - x^\ddagger)}{k_\mathrm{B}T}\right), \tag{S11b}$$

where $k_0$ is the pre-exponential factor, $x^\ddagger$ ($x_U - x^\ddagger$) are the relative distances between state $F$ ($U$) and the transition state. $\Delta G_{FU}$ is the free energy difference between states $F$ and $U$ at zero force. In a pulling experiment, the force is ramped linearly with time, $f = rt$, with $r$ the experimental pulling rate. The survival probability in the folded state (F) is

$$\frac{dP_F(f)}{df} = -\frac{k_{F\to U}(f)}{r} P_F(f)\,. \tag{S12}$$

By solving Eqs.(S11a), (S11b), and (S12) we get the unfolding rupture force distribution:

$$\begin{aligned} p_{F\to U}(f) = -\frac{dP_F(f)}{df} =& \frac{k_0}{r}\exp\left(\frac{k_0 k_{\rm B}T}{rx^\ddagger}\right)\exp\left(\frac{fx^\ddagger}{k_{\rm B}T}\right)\times \\ & \times\exp\left(-\frac{k_0 k_{\rm B}T}{rx^\ddagger}\exp\left(\frac{fx^\ddagger}{k_{\rm B}T}\right)\right) , \end{aligned} \tag{S13}$$

while for the reverse force distribution, one can similarly obtain

$$\begin{aligned} p_{U\to F}(f) =& \frac{\tilde{k}_0}{r}\exp\left(-\frac{f(x_U - x^\ddagger)}{k_{\rm B}T}\right)\times \\ & \times\exp\left(-\frac{\tilde{k}_0 k_{\rm B}T}{r(x_U - x^\ddagger)}\exp\left[-\frac{f(x_U - x^\ddagger)}{k_{\rm B}T}\right]\right) , \end{aligned} \tag{S14}$$

where we introduced $\tilde{k}_0 := k_0\exp(\Delta G_{FU}/k_{\rm B}T)$. We can recognize that the forward force distribution follows a Gompertz law, with an inverse scale parameter $s_U := k_{\rm B}T/x^\ddagger$ and with a mode given by $\mu_U = s_U\ln(r/k_0 s_U)$. This leads to the following useful re-parametrization:

$$p_{F\to U}(f) = \frac{1}{s_U}\exp\left(\frac{f-\mu_U}{s_U} + \exp\left(-\frac{\mu_U}{s_U}\right) - \exp\left(\frac{f-\mu_U}{s_U}\right)\right) . \tag{S15}$$

Eq.(S15) is very convenient as it is expressed in terms of the quantities $s_U, \mu_U$ whose order of magnitude can be easily estimated from experimental data (unlike $k_0$, $x^\ddagger$). For this reason, we used it both for MLE estimation (to retrieve $s_U$ and then $x^\ddagger$) and as a likelihood function in our Bayesian clustering algorithm (see Sec. S3).

The distance to the transition state $x^\ddagger$ is often expressed as a function of the variance of the rupture force distribution. Notice that there is no simple closed formula for $Var(X)$ when $X$ is a Gompertz-distributed random variable. For Gompertz distributions, the following approximation can be derived, $Var(X) \cong s_U^2\frac{\pi^2}{6}$ (72). It holds to a very good accuracy for the rupture force distributions measured in pulling experiments. As $\sqrt{\pi^2/6} \approx 1$, the rupture force distribution standard deviation is approximately equal to the inverse distance to the transition state per unit of $k_{\rm B}T$ in the BE model.

### S6 Continuous Effective Barrier Analysis

In the BE model, the height of the kinetic barrier decreases linearly with the applied force, $B(f) = B_0 - fx^\ddagger$. This hypothesis is relaxed in the kinetic diffusion (KD) model, which assumes the folding reaction as a diffusive process in a one-dimensional force-dependent free-energy landscape. The Continuous Effective Barrier Approach (CEBA) is based on the KD model. It can be used to extract the force-dependent behavior of the kinetic barrier from unzipping experiments (73, 74). In CEBA, the effective barrier between the folded ($F$) and the unfolded state ($U$), $B(f)$, is derived by imposing the detailed balance condition between the unfolding, $k_{FU}(f)$, and folding, $k_{FU}(f)$, kinetic rates (see Eqs.(S11)):

$$k_{F\to U}(f) = k_0 \exp\left(-\frac{B(f)}{k_\mathrm{B}T}\right) \tag{S16a}$$

$$k_{U\to F}(f) = k_{F\to U}(f) \exp\left(\frac{\Delta G_{FU}(f)}{k_\mathrm{B}T}\right), \tag{S16b}$$

where $k_0$ is the attempt rate, $B(f)$ is the effective barrier at force $f$, and $\Delta G_{FU}(f) = G_U(f) - G_F(f)$ is the folding free energy at force $f$. The latter term is given by

$$\Delta G_{FU}(f) = \Delta G_0 - \int_0^f \left(x_U(f') - x_F(f')\right) df', \tag{S17}$$

where $\Delta G_0$ is the folding free energy between $F$ and $U$ at zero force, and the integral accounts for the free energy change upon stretching the molecule in state $U$ ($F$) at force $f$.

We can derive two estimates for $B(f)$ by computing the logarithms of Eqs.(S16a) and (S16b), which give

$$\frac{B(f)}{k_\mathrm{B}T} = \log k_0 - \log k_{F\to U}(f) \tag{S18a}$$

$$\frac{B(f)}{k_\mathrm{B}T} = \log k_0 - \log k_{U\to F}(f) + \frac{\Delta G_{FU}(f)}{k_\mathrm{B}T}. \tag{S18b}$$

By imposing the continuity of the two estimations of $B(f)$ in Eqs.(S18), we can measure the folding free energy at force $f$, $\Delta G_{FU}(f)$. The free energy of the stretching contribution in $\Delta G_{FU}(f)$ Eq.(S17) can be measured from the unfolded branch. Matching Eqs.S18a and S18b permits us to directly estimate the folding free energy at zero force, $\Delta G_0$. Details can be found in (69, 74).

### S7 H1L12A thermodynamics for $\Delta C_p = 0$

A fit to the linear function $\Delta G_0 = \Delta H_0 - T\Delta S_0$ gives estimates for the folding enthalpy ($\Delta H_0$) and entropy ($\Delta S_0$) by assuming $\Delta C_p = 0$. We get $\Delta H_0^\mathrm{BAR} = 110(30)\,\mathrm{kcal\,mol^{-1}}$, $\Delta S_0^\mathrm{BAR} = 240(10)\,\mathrm{cal\,mol^{-1}K^{-1}}$ (continuous blue line in Fig. S9) and $\Delta H_0^\mathrm{CEBA} = 100(30)\,\mathrm{kcal\,mol^{-1}}$, $\Delta S_0^\mathrm{CEBA} = 230(8)\,\mathrm{cal\,mol^{-1}K^{-1}}$ (continuous red line in Fig. S9). Our results for $\Delta G_0$ agree with the Mfold prediction at 37°C (black triangles in Fig. S9) above room temperature. However, a discrepancy is observed for the enthalpy and entropy values, $\Delta H_0^\mathrm{Mfold} = 196\,\mathrm{kcal\,mol^{-1}}$, $\Delta S_0^\mathrm{Mfold} = 533\,\mathrm{cal\,mol^{-1}K^{-1}}$, almost twice our numbers. In contrast, by assuming $\Delta C_p \neq 0$ and applying the Clausius-Clapeyron equation (see main text), we measured

$\Delta H_0^{\mathrm{BAR}} = 163(6)\,\mathrm{kcal\,mol^{-1}}$ and $\Delta S_0^{\mathrm{BAR}} = 420(20)\,\mathrm{cal\,mol^{-1}K^{-1}}$ at 37°C. In this case, we only report results obtained from $\Delta G_0$ measurements derived with the BAR method, available at all $T$ for N and at 7°C for M.

### S8 H1L4A and H1L12A thermodynamics for $\Delta C_p \neq 0$

We have measured the enthalpy $\Delta H_0$ and entropy $\Delta S_0$ of N at all temperatures for hairpins H1L4A and H1L12A using the Clausius-Clapeyron equation under an applied force (61). Both $\Delta H_0$ and $\Delta S_0$ are temperature dependent indicating a finite $\Delta C_p = \frac{\partial \Delta H}{\partial T} = T\frac{\partial \Delta S}{\partial T}$. We observe two distinct regimes, above (hot, H) and below (cold, C) $\sim$ 20°C (see Fig. 5B, main text). By separately fitting the two regimes to the function $\Delta S_0(T) = \Delta S_{\mathrm{H(C)}} + \Delta C_p \log(T/T_{\mathrm{H(C)}})$ where $\Delta S_{\mathrm{H(C)}}$ is the entropy at the hot (cold) denaturation temperature $T_{\mathrm{H(C}}$, we find $\Delta C_p^{\mathrm{H}} = 1.5(2) \cdot 10^3$ cal mol$^{-1}$K$^{-1}$ and $\Delta C_p^{\mathrm{C}} = 5.8(4) \cdot 10^3$ cal mol$^{-1}$K$^{-1}$ for H1L4A, and $\Delta C_p^{\mathrm{H}} = 1.5(2) \cdot 10^3$ cal mol$^{-1}$K$^{-1}$ and $\Delta C_p^{\mathrm{C}} = 8(1) \cdot 10^3$ cal mol$^{-1}$K$^{-1}$ for H1L12A. Notice that by measuring $\Delta C_p^{\mathrm{H(C)}}$ from the enthalpy $T$-dependence, we obtain compatible results: $\Delta C_p^{\mathrm{H}} = 1.3(2) \cdot 10^3$ cal mol$^{-1}$K$^{-1}$ and $\Delta C_p^{\mathrm{C}} = 6.0(4) \cdot 10^3$ cal mol$^{-1}$K$^{-1}$ for H1L4A, and $\Delta C_p^{\mathrm{H}} = 1.7(3) \cdot 10^3$ cal mol$^{-1}$K$^{-1}$ and $\Delta C_p^{\mathrm{C}} = 7(1) \cdot 10^3$ cal mol$^{-1}$K$^{-1}$ for H1L12A.

$\Delta C_p$ values permit us to determine hot and cold denaturation temperatures, which are $T_{\mathrm{H}} = 377(3)$K and $T_{\mathrm{C}} = 222(3)$K for H1L4A, and $T_{\mathrm{H}} = 366(4)$K and $T_{\mathrm{C}} = 228(4)$K for H1L12A. Despite the differences in $\Delta C_p^{\mathrm{C}}$ between H1L4A and H1L12A, in both cases, stability is maximum at $T_S \sim 5$°C where $\Delta S_0 = 0$. Moreover, cold denaturation is predicted at $T_C \sim -50(4)$°C for both sequences. This concurrency suggests that maximum stability and cold denaturation temperatures are sequence-independent.

Finally, we report the entropy at the hot and cold denaturation temperatures from the fit $\Delta S_{\mathrm{H}} = 630(40)$ cal mol$^{-1}$K$^{-1}$ and $\Delta S_{\mathrm{C}} = -1340(90)$ cal mol$^{-1}$K$^{-1}$ for H1L4A, and $\Delta S_{\mathrm{H}} = 670(40)$ cal mol$^{-1}$K$^{-1}$ and $\Delta S_{\mathrm{C}} = -1600(200)$ cal mol$^{-1}$K$^{-1}$ for H1L12A.

### Supplementary Figures

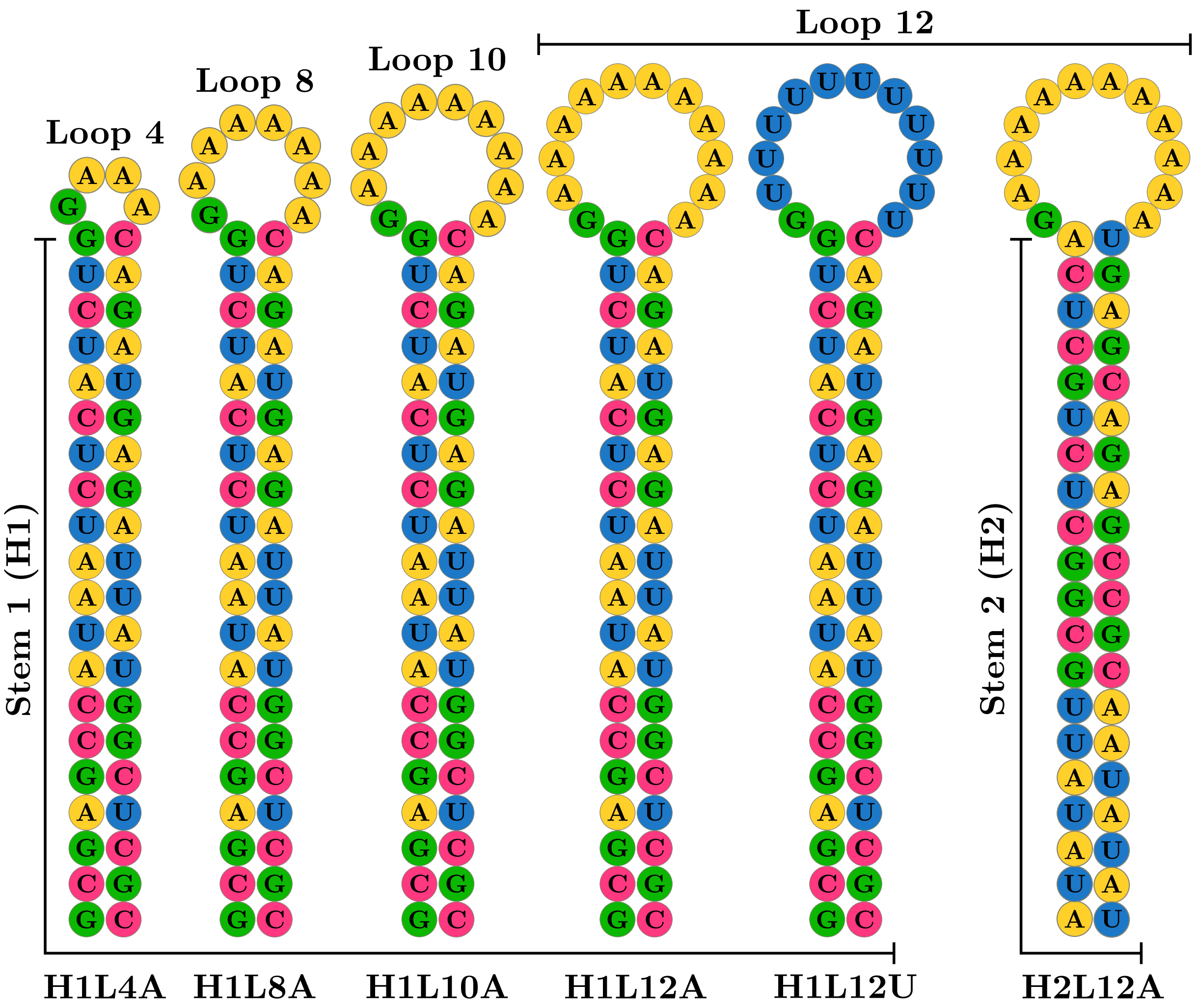

Figure S1: **Sequence of the studied RNA hairpins.** Each hairpin is named after the stem sequence (H1 or H2), the loop size (L4, L8, L10, or L12), and the loop composition (poly-A or poly-U). The sequences are reported in Table S1

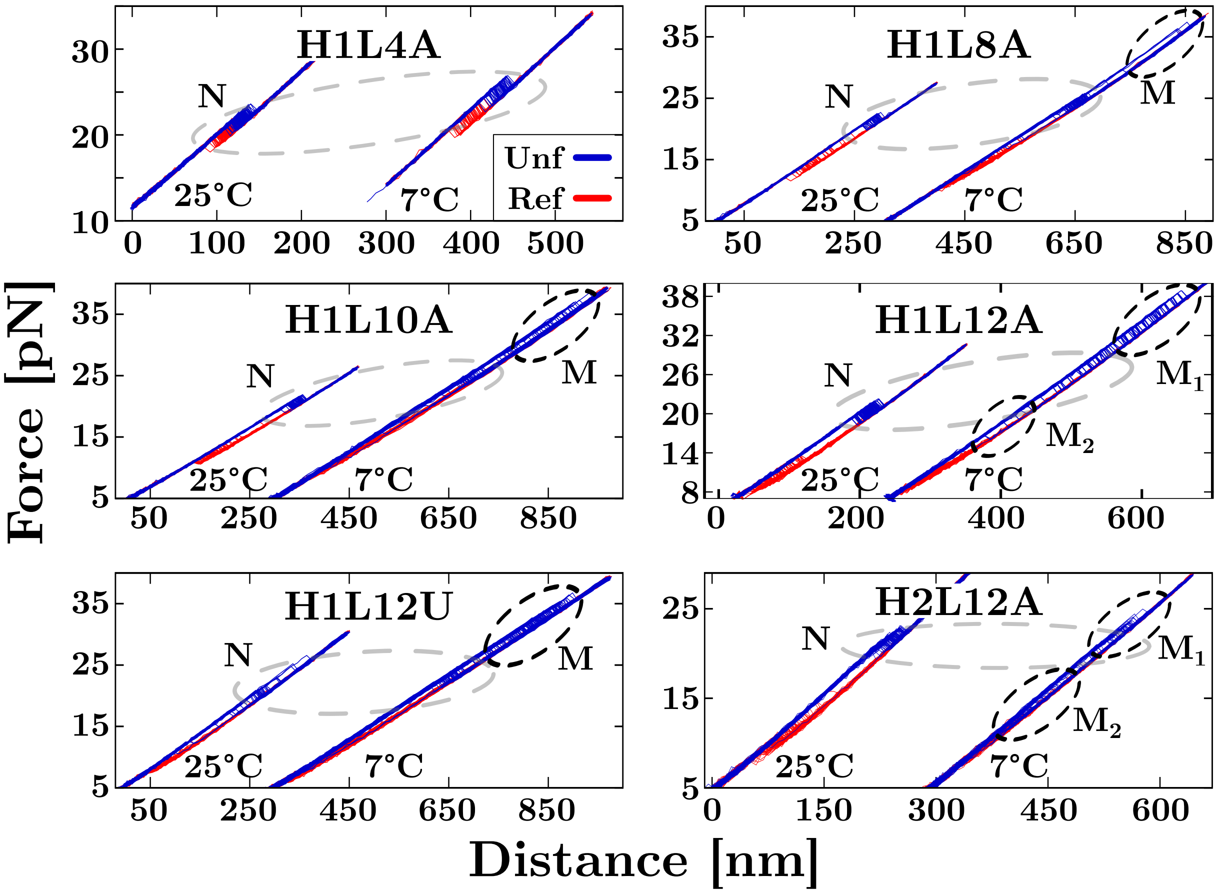

Figure S2: **RNA unzipping experiments in sodium.** Experimental FDCs (25°C and 7°C) measured at 1M NaCl for all the studied hairpins. At low-$T$, all molecules exhibit analogous behavior to that observed at 4mM $MgCl_2$.

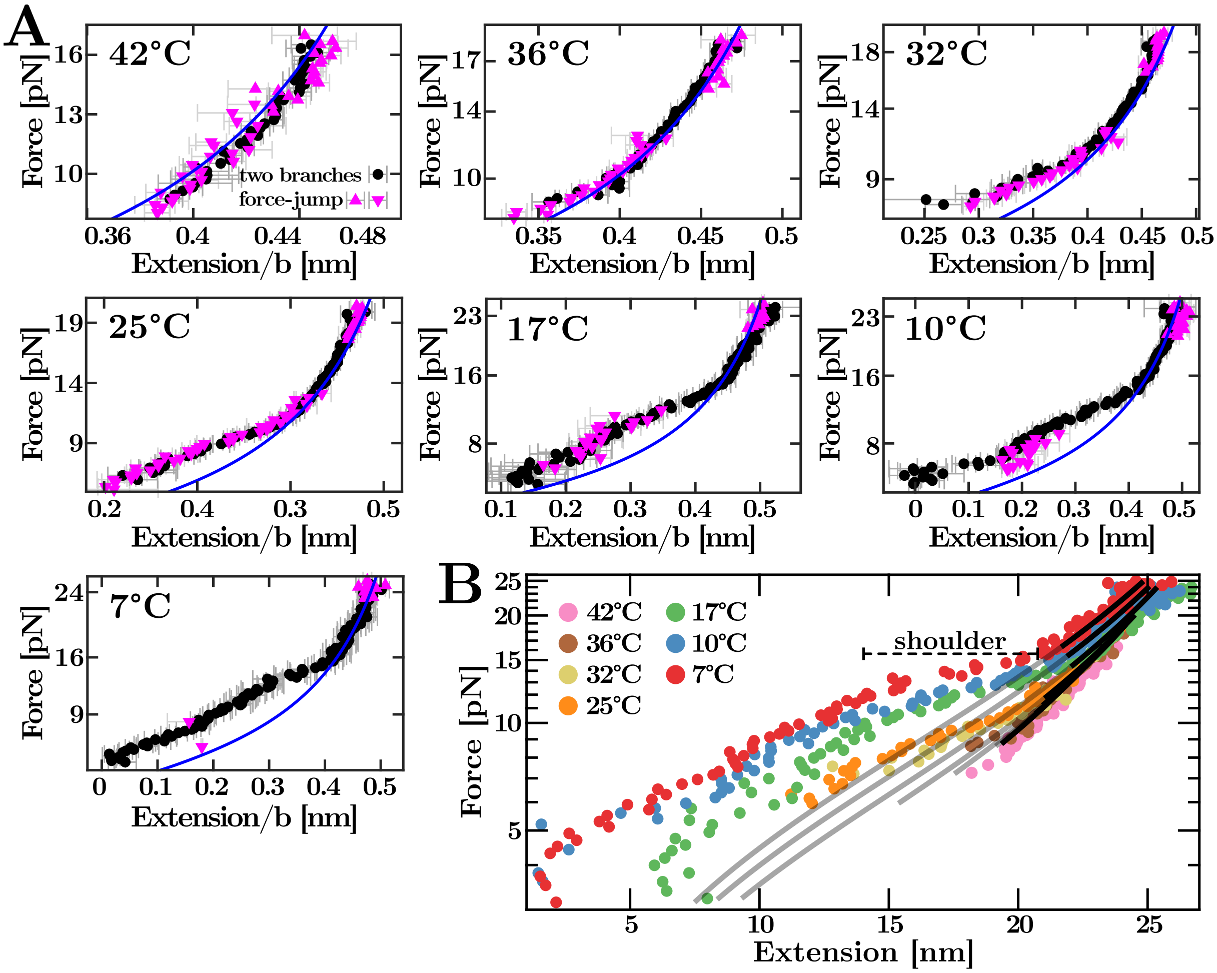

Figure S3: $T$**-dependent H1L12A ssRNA elastic response.** (**A**) Force versus the ssRNA extension per base at all the studied temperatures. Two methods have been used to extract the molecular extension of the ssRNA: the force-jump (magenta triangles up -unfolding- and down -refolding-) and the two-branches method (black circles) (58, 59). Blue lines are the fits to the WLC.(**B**) Overview of the $T$-dependent ssRNA elastic response. Only data above the shoulder (horizontal-dashed segment) have been used for the fit (black lines). Grey lines extrapolate the WLC fitting curves to the non-fitting regions.

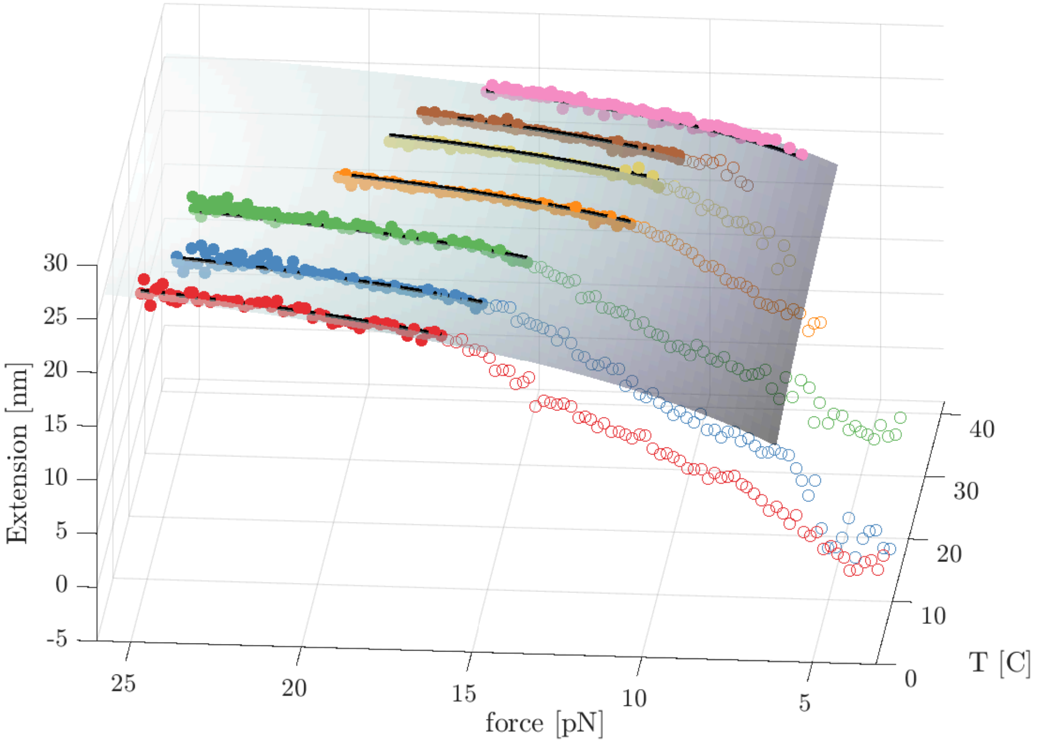

Figure S4: **Multi-$T$ fit of the H1L12A ssRNA elastic response**. We simultaneously fit the relation $l_p = l_p(T) = a_1 T + b_1$, $l_B = l_B(T) = a_2 T + b_2$ on data points at all temperatures. This gives for $l_p$ the values $a_1 = 0.135 \pm 0.006$ Å/C, $b_1 = 3.5 \pm 0.1$ Å and for $l_B$ the values $a_2 = -0.023 \pm 0.002$ Å/C, $b_2 = 7.37 \pm 0.05$ Å. The fit is performed over the filled symbols only. The three-dimensional $T$-force-extension surface is represented in light grey. The black lines plot force-extension cross-sections at a given temperature (red, $T = 7$°C; blue, $T = 10$°C; green, $T = 17$°C; orange, $T = 25$°C; yellow, $T = 32$°C; brown, $T = 36$°C; pink, $T = 42$°C).

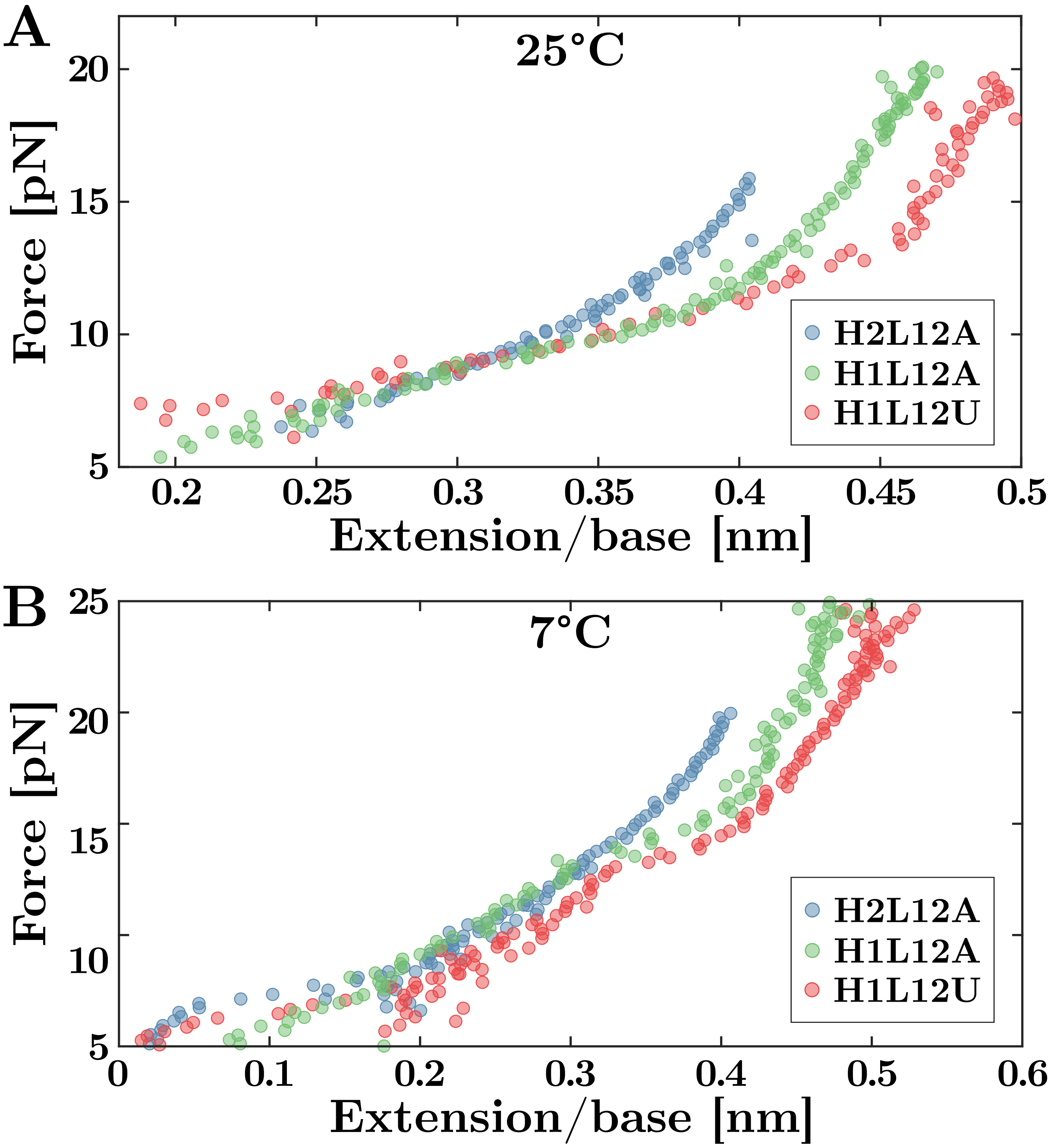

Figure S5: **$T$-dependence of the H1L12A, H2L12A, and H1L12U ssRNA elastic response.** Results are shown at $T = 25°C$ (panel A) and 7°C (panel B).. Extension is normalized per base by dividing the measured extension by each hairpin's total number of bases. The normalized extensions do not collapse as different sequences feature different elastic properties (see main text).

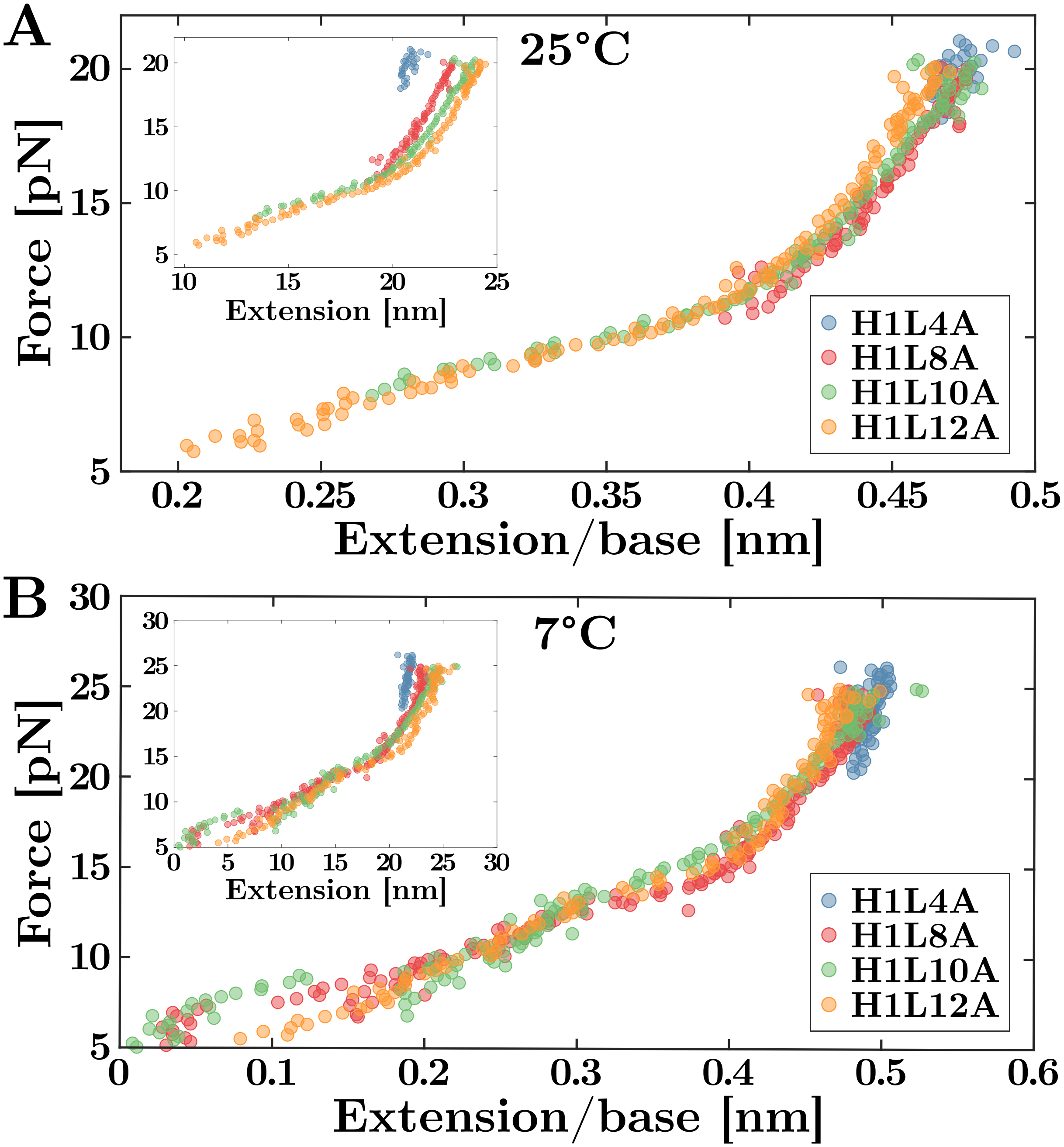

Figure S6: $T$**-dependent ssRNA response for H1L4A, H1L8A, H1L10A, and H1L12A.** Results are shown at $T = 25°\text{C}$ (panel A) and $7°\text{C}$ (panel B). If we plot the force versus the total extension, data for different hairpins do not collapse (insets of A and B). In contrast, upon normalizing the extension per base, the force-extension curves of all hairpins collapse into a master curve (main A and B). Extension is normalized per base by dividing the measured extension by each hairpin's total number of bases.

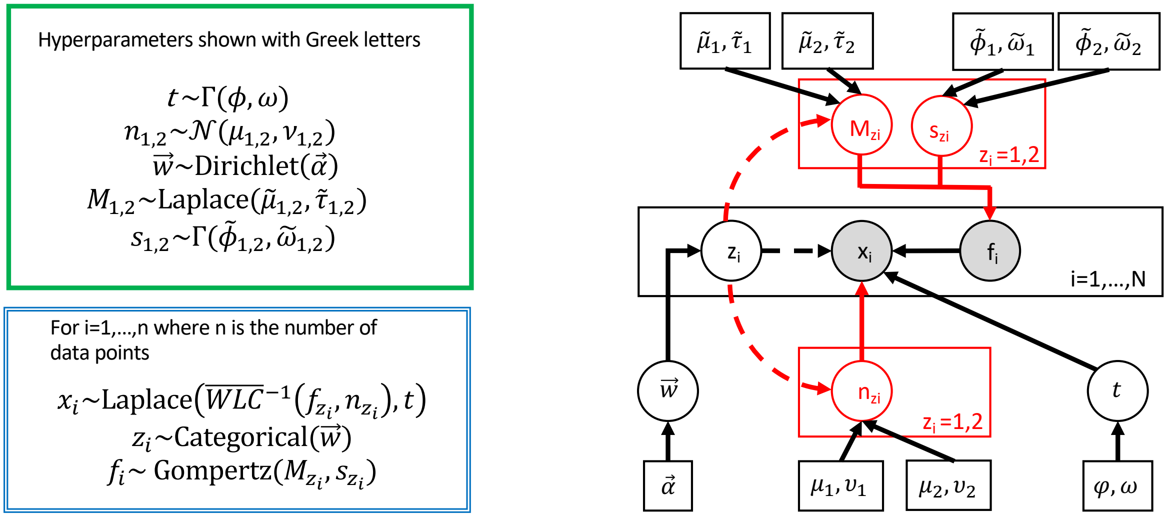

Figure S7: **The Bayesian classification algorithm.** (**Left**) Specification of the prior (in green) and likelihood (in blue) functions used. The hyper-parameters used are indicated by Greek letters. $\Gamma$ stands for the gamma distribution, $\mathcal{N}$ for the normal distribution. (**Right**) Probabilistic graph view of the Bayesian network used. Misfolded and native states are represented by the superscript $1, 2$ and are encoded in the latent variables $z_i = 1, 2$. We highlight in red the part of the model that depends on $z_i$: the rupture force distribution through its mode $M_{z_i}$ and scale $1/s_{z_i}$ parameters and the number of monomers released within an unfolding event $n_{z_i}$.

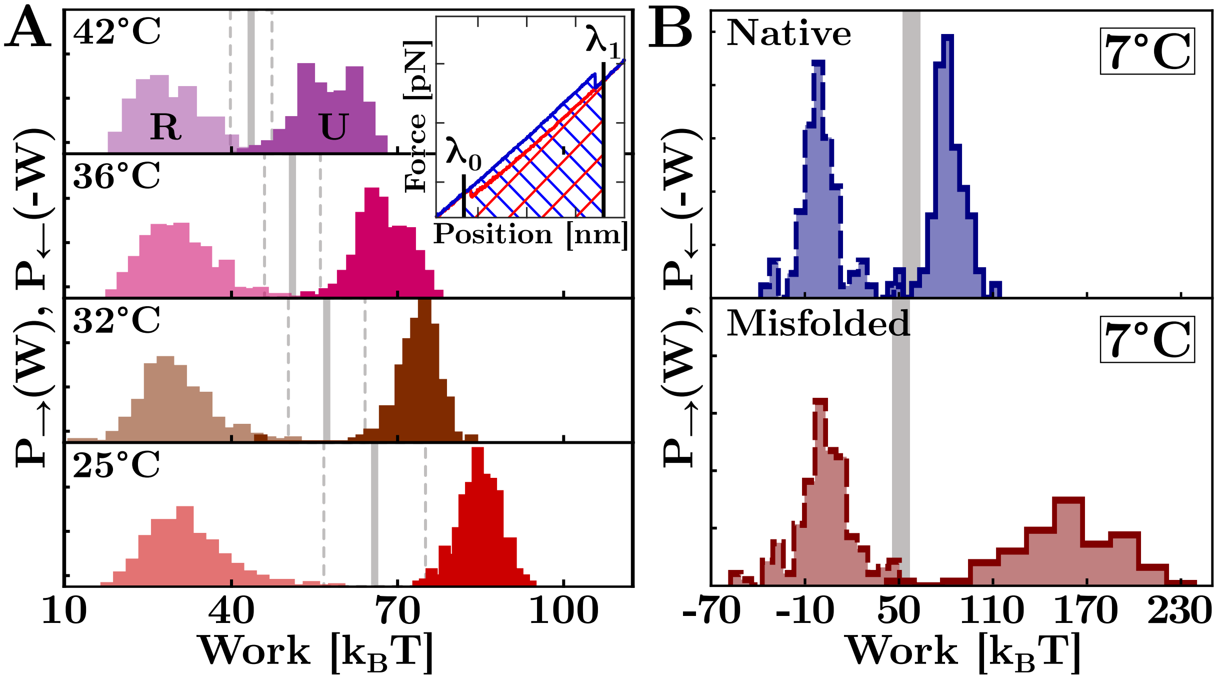

Figure S8: **Fluctuation theorem applied on the H1L12A.** (**A**) Unfolding (U) and refolding (R) work distributions, and $\Delta G_0$ estimates (gray thick lines) at 4mM $MgCl_2$ above 25°C. Results are averages over 5-6 molecules. Dashed grey lines show the hysteresis region. **Inset.** The measured work equals the area under the FDC for unfolding (blue) and refolding (red). (**B**) $P_{\rightarrow}(W)$ (solid line) and $P_{\leftarrow}(-W)$ (dashed line) for N (blue) and M (red) states at 4mM $MgCl_2$ and 7°C.

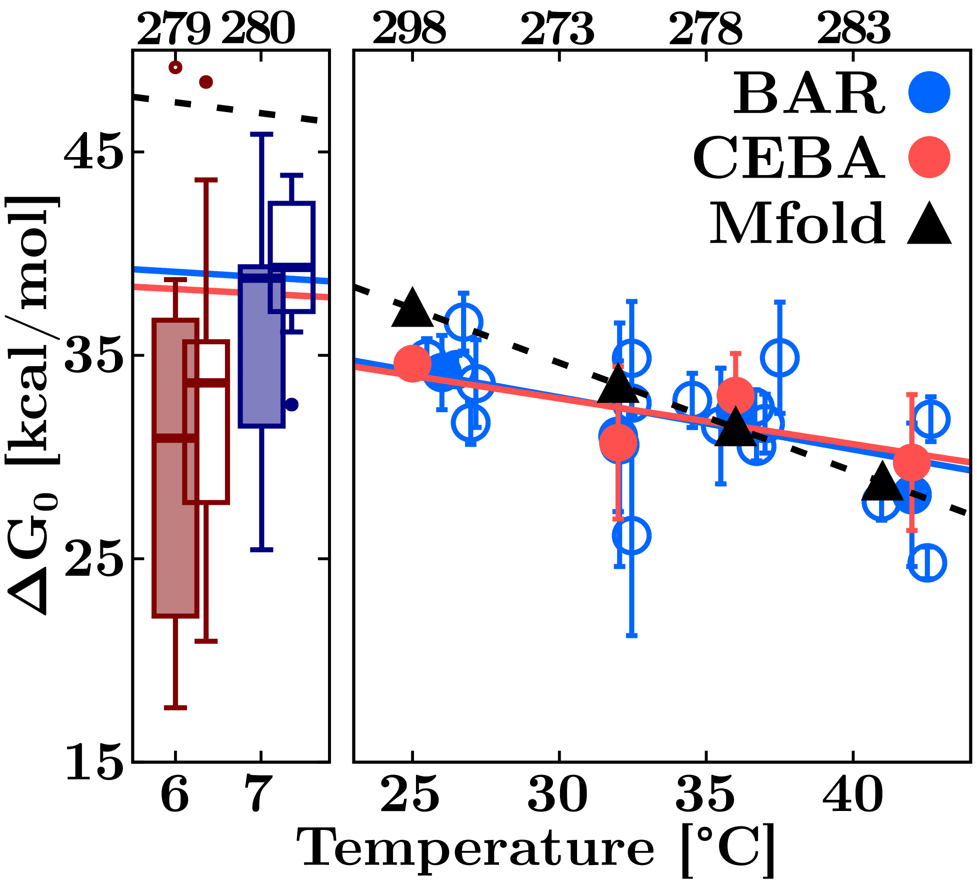

Figure S9: $T$**-dependence of the H1L12A free energy for N and M**. $\Delta G_0$ values at 7°C in 4mM $MgCl_2$ (solid boxes) and 400mM NaCl (empty boxes) **(left)**, and above 25°C at 4mM $MgCl_2$ for N **(right)**. **(Left)** Results are shown with a box-and-whisker plot indicating the data median (horizontal thick line), first and third quartiles (box), 10th and 90th percentiles (whiskers), and outliers (dots). **(Right)** BAR (blue) and CEBA (red) results are compared with Mfold prediction (black). For BAR, we show results for different molecules (empty circles) and their averages (solid circles). Temperature axis in °C (bottom label) and K (top label).

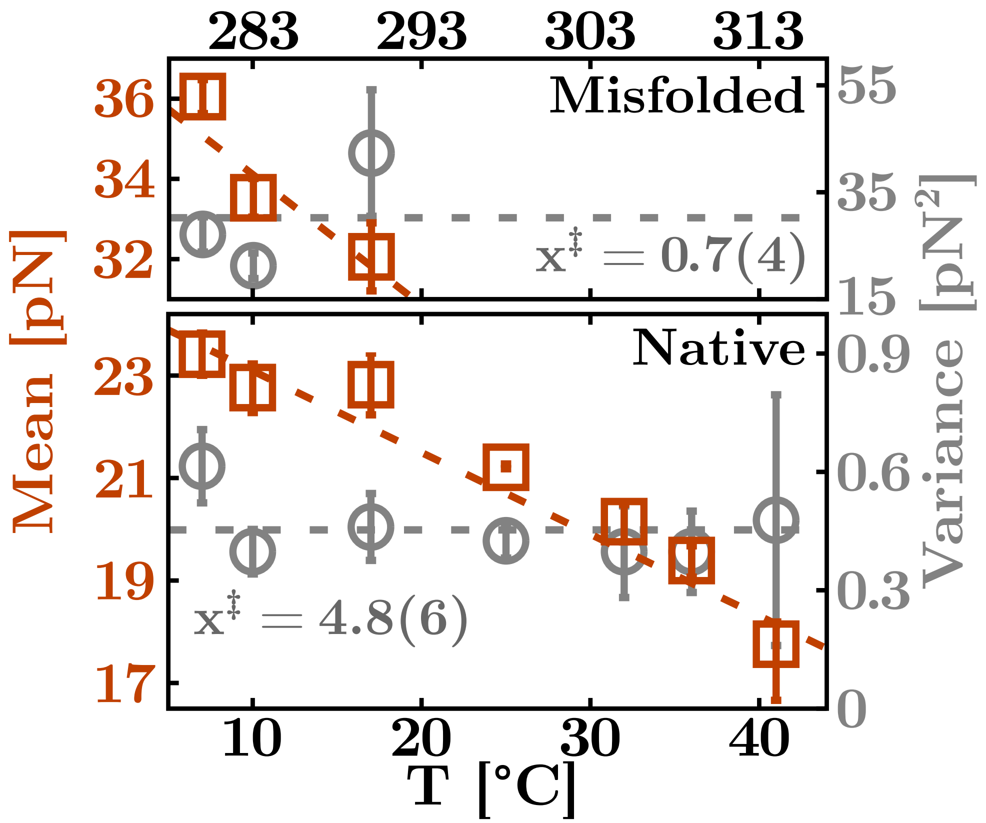

Figure S10: **Transition state distance in H1L12A.** Mean (orange squares) and variance (grey circles) of the rupture force distributions for native (bottom) and misfolded (top) at each experimental $T$ (in Celsius and Kelvin) for H1L12A. The dashed grey lines denote the average variance used to measure the transition state distances for N and M.

### Supplementary Tables

| Molecule | Stem Sequence | Loop Sequence |
|---|---|---|
| H1L4A | GCGAGCCAUAAUCUCAUCUG | GAAA |
| H1L8A | GCGAGCCAUAAUCUCAUCUG | GAAAAAAA |
| H1L10A | GCGAGCCAUAAUCUCAUCUG | GAAAAAAAAA |
| H1L12A | GCGAGCCAUAAUCUCAUCUG | GAAAAAAAAAAA |
| H1L12U | GCGAGCCAUAAUCUCAUCUG | GUUUUUUUUUUU |
| H2L12A | AUAUAUUGCGGCUCUGCUCA | GAAAAAAAAAAA |

Table S1: Sequences of the six RNA hairpins ($5' \rightarrow 3'$, from left to right).

| | | H1L12A | | H1L12U | | H2L12A | |
|---|---|---|---|---|---|---|---|
| **T [°C]** | **T [K]** | $\mathbf{l_p}$ | $\mathbf{d_b}$ | $\mathbf{l_p}$ | $\mathbf{d_b}$ | $\mathbf{l_p}$ | $\mathbf{d_b}$ |
| 42 | 315 | 0.99(3) | 0.64(1) | | | | |
| 36 | 309 | 0.81(2) | 0.66(1) | | | | |
| 32 | 305 | 0.79(3) | 0.65(1) | | | | |
| 25 | 298 | 0.70(2) | 0.67(1) | 0.60(3) | 0.75(1) | 0.55(3) | 0.67(2) |
| 17 | 290 | 0.46(2) | 0.76(1) | | | | |
| 10 | 283 | 0.42(3) | 0.75(2) | | | | |
| 7 | 280 | 0.38(3) | 0.75(2) | 0.35(3) | 0.82(2) | 0.28(2) | 0.77(2) |

Table S2: Temperature dependence of the persistence length ($l_p$) [nm] and the interphosphate distance ($d_b$) [nm] of the different dodecaloop hairpins. The error (in brackets) refers to the last digit.

| T [°C] | T [K] | BAR | CEBA | Mfold |
|---|---|---|---|---|
| 42 | 315 | 28 (3) | 30 (3) | 28 |
| 36 | 309 | 32 (1) | 33 (2) | 31 |
| 32 | 305 | 31 (4) | 31 (4) | 34 |
| 25 | 298 | 34 (2) | 35 (1) | 37 |

Table S3: $\Delta G_0$ [kcal/mol] values of N for $\mathrm{H1L12A}$ in the high-$T$ regime measured with BAR and CEBA methods compared to predictions by Mfold. The error (in brackets) refers to the last digit.

| **T [°C]** | **T [K]** | **State** | **$Mg^{2+}$** | **$Na^{+}$** |
|---|---|---|---|---|
| 7 | 280 | N | 38 (9) | 37 (3) |
| | | M | 31 (10) | 31 (8) |

Table S4: $\Delta G_0$ [kcal/mol] values of M for $\mathrm{H1L12A}$ at $T = 7$°C derived with the BAR method. To be compared, the $100/1$ equivalence rule between monovalent and divalent salt concentrations has been applied to the sodium results. The error (in brackets) refers to the last digit.

| T [°C] | T [K] | $\Delta$H [kcal mol$^{-1}$] | $\Delta$S [cal mol$^{-1}$ K$^{-1}$] |
|---|---|---|---|
| 42 | 315 | 165 (7) | 433 (18) |
| 36 | 309 | 163 (6) | 421 (19) |
| 32 | 305 | 152 (7) | 396 (18) |
| 25 | 298 | 139 (5) | 351 (17) |
| 17 | 290 | 114 (8) | 263 (15) |
| 10 | 283 | 71 (6) | 110 (9) |
| 7 | 280 | 39 (10) | 2 (3) |

Table S5: Temperature dependence of the enthalpy ($\Delta H$) and entropy ($\Delta S$) of N for H1L12A. The error (in brackets) refers to the last digit(s).

| **T [°C]** | **T [K]** | $\mathbf{\Delta G_0}$ [$\mathbf{kcal\,mol^{-1}}$] | $\mathbf{\Delta H}$ [$\mathbf{kcal\,mol^{-1}}$] | $\mathbf{\Delta S}$ [$\mathbf{cal\,mol^{-1}\,K^{-1}}$] |
|---|---|---|---|---|
| 40 | 313 | 30 (3) | 345 (35) | 138 (11) |
| 32 | 305 | 33 (4) | 316 (34) | 130 (11) |
| 25 | 298 | 37 (3) | 269 (33) | 117 (10) |
| 15 | 288 | 39 (5) | 155 (27) | 84 (9) |
| 10 | 283 | 38 (4) | 61 (19) | 56 (7) |
| 7 | 280 | 38 (4) | -12 (12) | 34 (5) |

Table S6: Temperature dependence of the free energy ($\Delta G_0$), enthalpy ($\Delta H$), and entropy ($\Delta S$) of N for H1L4A. The error (in brackets) refers to the last digit(s).
